# Immune cell multi-omics analysis reveals contribution of oxidative phosphorylation to B cell functions and organ damage of lupus

**DOI:** 10.1101/2021.10.08.463629

**Authors:** Yusuke Takeshima, Yukiko Iwasaki, Masahiro Nakano, Yuta Narushima, Mineto Ota, Yasuo Nagafuchi, Shuji Sumitomo, Tomohisa Okamura, Keith B Elkon, Kazuyoshi Ishigaki, Akari Suzuki, Yuta Kochi, Kazuhiko Yamamoto, Keishi Fujio

## Abstract

**Objective:** Systemic lupus erythematosus (SLE) is the prototypical systemic autoimmune disease, with a poor long-term prognosis. The type I interferon (IFN) signature, a prominent feature of SLE, is not an ideal therapeutic target or outcome predictor. To explore immunological pathways in SLE more precisely, we performed integrative analysis of transcriptomics, epigenomics, and genomics using each immune cell subset from peripheral blood.

**Methods:** We sorted 18 immune cell subsets and identified the mRNA expression profiles and genetic polymorphisms in 107 SLE patients and 92 healthy controls. Open chromatin information was also taken by ATAC-seq analysis. Combined differentially expressed genes (DEGs) and expression quantitative trait loci (eQTL) analysis was conducted to find key driver genes in SLE pathogenesis.

**Results:** We found transcriptomic, epigenetic, and genetic importance of oxidative phosphorylation (OXPHOS)/mitochondrial dysfunction in SLE memory B cells. Particularly, we identified an OXPHOS-regulating gene, *PRDX6*, as a key driver in SLE B cells. *Prdx6*–deficient B cells showed upregulated mitochondrial respiration as well as antibody production. We revealed OXPHOS signature was associated with type I IFN signaling-related genes (ISRGs) signature in SLE memory B cells. Furthermore, the gene sets related to innate immune signaling among ISRGs presented correlation with OXPHOS and these two signatures showed associations with SLE organ damage as well as specific clinical phenotypes.

**Conclusion:** This work elucidated the potential prognostic marker for SLE. Since OXPHOS consists of the electron transport chain, a functional unit in mitochondria, these findings suggest the importance of mitochondrial dysfunction as a key immunological pathway involved in SLE.

## INTRODUCTION

Systemic lupus erythematosus (SLE), which is a prototypic systemic autoimmune disease, exhibits a poor long-term prognosis due to the accumulation of organ damage. Previous studies on SLE, including genome-wide association studies (GWAS) and gene expression studies in peripheral blood mononuclear cells (PBMCs), indicated a role of type I interferon (IFN) signaling in SLE immunological pathogenesis [1–3]. However, the type I IFN signature does not correlate with long-term prognosis [4–9], and, although Phase III trials with an antibody blocking the type I IFN pathway showed improvement of clinical activity, approximately 50% of SLE patients did not exhibit a significant response [10]. These observations strongly suggest a pathogenic contribution from critical immunological pathways other than the classical type I IFN signature. Consistently, based on recent progress in integrated analyses, researchers have revealed a correlation between lupus nephritis activity and a plasmablast/neutrophil signature in blood [11]. Moreover, the exhausted CD8 T cell gene signature is associated with good outcomes [9]. In order to advance these findings, more detailed analyses of immune cell subsets are needed because transcriptome analyses of a mixture of cell subsets are obscured by variations among the different cell types. Moreover, the subset-derived transcriptome has less variability than the single-cell-derived transcriptome, and suitable for association analysis with genetic polymorphisms with higher accuracy.

In this study, we analyzed the transcriptomes of 18 blood immune cell subsets and the genotypes of 107 SLE patients and 92 healthy controls (HCs), together with open chromatin data obtained by Assay for Transposase-Accessible Chromatin (ATAC)-seq. Our integrated analysis indicated the importance of B cell metabolic regulation via mitochondrial function in SLE pathogenesis. In addition, we identified key driver genes (KDGs) related to mitochondrial dysfunction in the SLE B cells using eQTL analysis. Notably, gene sets related to innate immune signaling, including an oxidative phosphorylation (OXPHOS) signature, showed associations with SLE organ damage.

## RESULTS

### The importance of memory B cells in SLE pathogenesis via OXPHOS/mitochondrial dysfunction according to open chromatin and transcriptome analyses

We performed an integrated analysis using the study pipeline shown in figure 1A and data quality was checked as in online supplemental figure S1A. We first evaluated the relative genetic contribution of each immune cell subset in PBMCs to SLE pathogenesis. We applied stratified linkage disequilibrium (LD) score regression (LDSC) [13] for partitioning heritability to identify SLE-relevant cell types using our ATAC-seq data sets. We identified an enrichment of genes related to SLE risk within the open chromatin of B cells, especially in three memory B cell subsets: unswitched memory B cells (USM B), switched memory B cells (SM B), and double-negative B cells (DN B) (figure 1B). This was consistent with a previous study that demonstrated an enrichment of SLE risk in the open chromatin of whole B cells [14].

**Figure 1.**
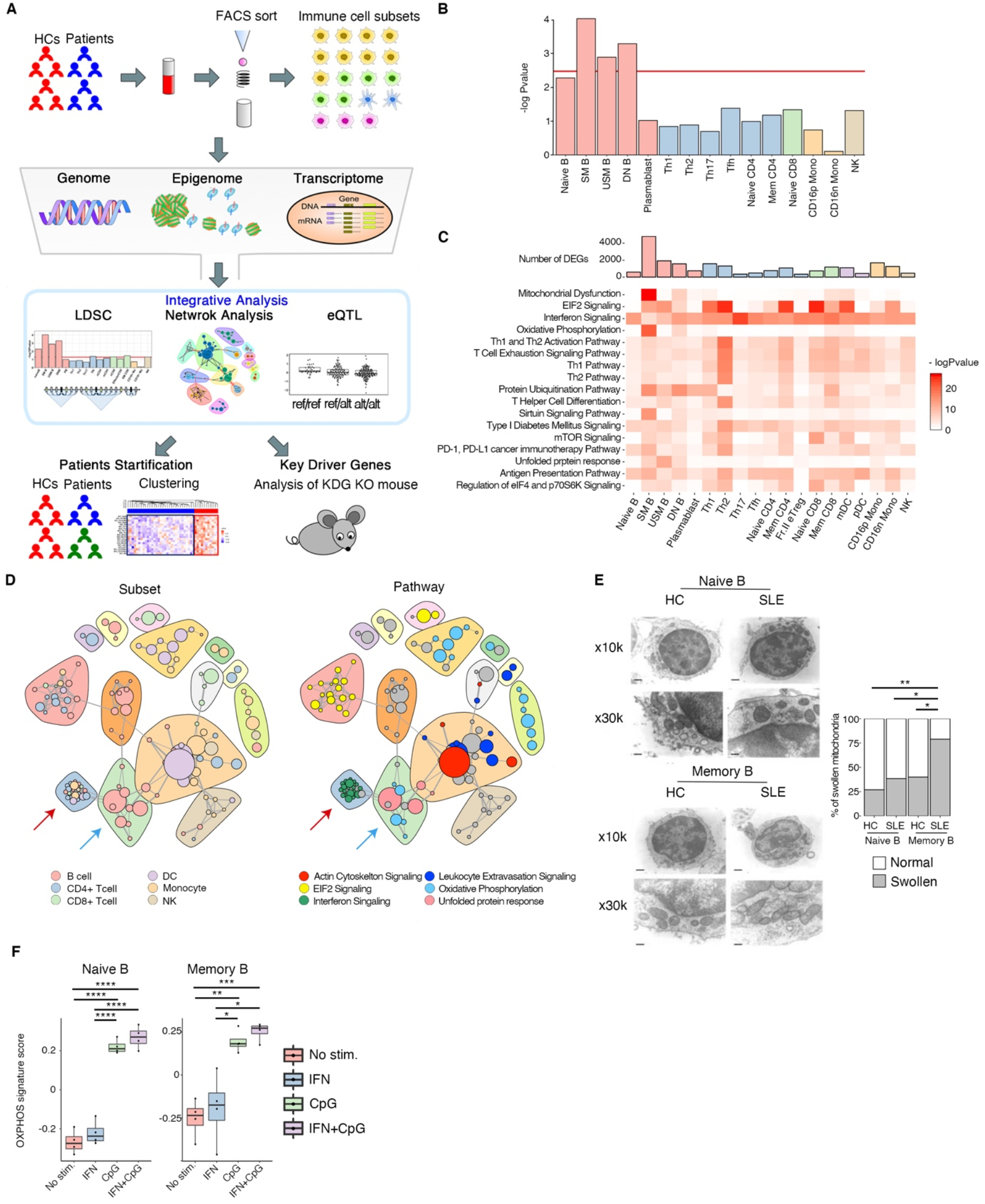
Overview of our analysis and identification of mitochondrial function as a key on SLE memory B cells. (**A**) Schematic view of the work in this study. Pipeline for collecting the 19 immune cell subsets from PBMCs, and the genomic, epigenomic, and transcriptomic data for the subsequent analysis. (**B**) Linkage score regression analysis of open chromatin data using summary statistics from a GWAS of SLE patients. Red line indicates significance at *p* < 0.05 after Bonferroni correction for the 15 immune cell types tested. (**C**) Pathway analysis of DEGs in all immune cell subsets with *q* < 0.05 in the test cohort. The pathways with –log10 *p*-values > 10 in any immune cell subset were visualized by heatmaps. (**D**) Network analysis of each module correlation in the test cohort. The top three pathways with high –log10 *p*-values that were annotated in at least two modules by IPA were selected. Only the modules with *r*^2^ > 0.7 were visualized. The line thickness reflects the correlation strength. Red arrow indicates IFN signaling modules. Blue arrow indicates OXPHOS modules in memory B cells. (**E**) Transmission electron microscopy of purified naïve B and memory B cells from HCs and SLE patients. Percentages of cells with swollen mitochondria (diameter > 500 nm) among 20 analyzed cells are shown. (**F**) Human naïve and memory B cells were cultured for 72h with combinations of CpG ODN 2006 and IFN-α. CpG ODN 2006 (2.5 μg/ml) and recombinant human IFN-α (1000 U/ml) were used. OXPHOS signature score was calculated in each condition. n = 4. Student’s t test was performed. * *p* < 0.05, ** *p* < 0.01, *** *p* < 0.001, **** *p* < 0.0001.

Our transcriptomic data showed widespread perturbations in SLE-related genes, with approximately 1,000 differentially expressed genes (DEGs) detected in each immune cell subset. We also found an enrichment of SLE risk within these DEGs by LDSC analysis of specifically expressed genes (online supplemental figure S1B). Pathway analysis of the DEGs revealed a conserved IFN signaling pathway among all immune cell subsets. With regard to B cells, we revealed a relative enrichment of DEGs related to OXPHOS and mitochondrial dysfunction pathways (figure 1C and online supplemental figure S1C). Due to an almost complete overlap of genes between the OXPHOS and mitochondrial dysfunction pathways, we used the term “OXPHOS” in our following analyses. Using weighted gene correlation network analysis (WGCNA) [15], we determined the correlations of each module eigengene among all subsets in the test and validation cohorts (figure 1D and online supplemental figure S1D). IFN signaling modules showed a strong correlation with each other irrespective of the immune cell type. Interestingly, OXPHOS modules in USM B and SM B were correlated with these IFN signaling modules more strongly than in other cell types. This result suggests a biological relationship between these two pathways in memory B cells, consistent with previous reports showing the importance of OXPHOS in SLE [16,17]. Furthermore, transmission electron microscopy revealed an increased proportion of swollen mitochondria [18] in SLE memory B cells, but not SLE naive B cells (figure 1E). The upregulated mitochondria-related genes did not suggest apoptosis of memory B cells in SLE patients, because neither naive nor memory B cells in SLE patients were pro-apoptotic (online supplemental figure S1E) [19]. Differentiation of memory B cells into plasmablasts by stimulation with a Toll-like receptor (TLR) 9 agonist (CpG) and type I IFN [20,21] was inhibited by administration of inhibitors of the electron transport chain (ETC) complex I or III (online supplemental figure S1F), confirming the importance of OXPHOS in plasmablast differentiation. Notably, in stimulated memory B cells, OXPHOS signature scores were induced by CpG, but not type I IFN (figure 1F), suggesting a role for innate immune signaling for inducing the OXPHOS pathway in SLE B cells.

### Epigenetic regulation of the OXPHOS signature according to ATAC-seq

Next, we evaluated whether upregulation of the OXPHOS signature is associated with an open chromatin state, using ATAC-seq data. Consistent with the LDSC results in figure 1B, differentially accessible regions (DARs) were most abundant in memory B cells (figure 2A). Next, we applied the chromVAR algorithm to our ATAC-seq data sets to investigate cell type-specific transcriptional regulation with accessibility of transcription factor (TF). [22]. Firstly, principal component analysis showed each parental immune cell type was clustered independently (figure 2B), supporting the quality of our ATAC-seq analysis. Each B cell subset was clustered independently, and disease status constituted distinct clusters within each B cell subset (figure 2C). Because the expression of ETC genes was upregulated in all B cell subsets in SLE patients (online supplemental figure S2A and online supplemental table S1), we evaluated the open chromatin status within the binding regions of TFs that regulate the ETC complex, including nuclear respiratory factor 1 and 2 (NRF1 and NRF2), estrogen-related receptor alpha, yin yang 1 (YY1), and CREB [23,24]. The TF enrichment scores for NRF1, YY1, estrogen response element (ERE), and cAMP response element (CRE) showed a higher tendency to be present in each B cell subset of SLE patients (figure 2D). The enrichment scores of YY1 in plasmablast and CRE in Naive B cells and plasmablasts showed a significant upregulation (figure 2E). Moreover, CREB1, which is considered to interact with CRE sites in mtDNA, thereby regulating ETC expression [25], showed significant high enrichment score for Naive B cells and plasmablasts in SLE patients (online supplemental figure S2B).

**Figure 2.**
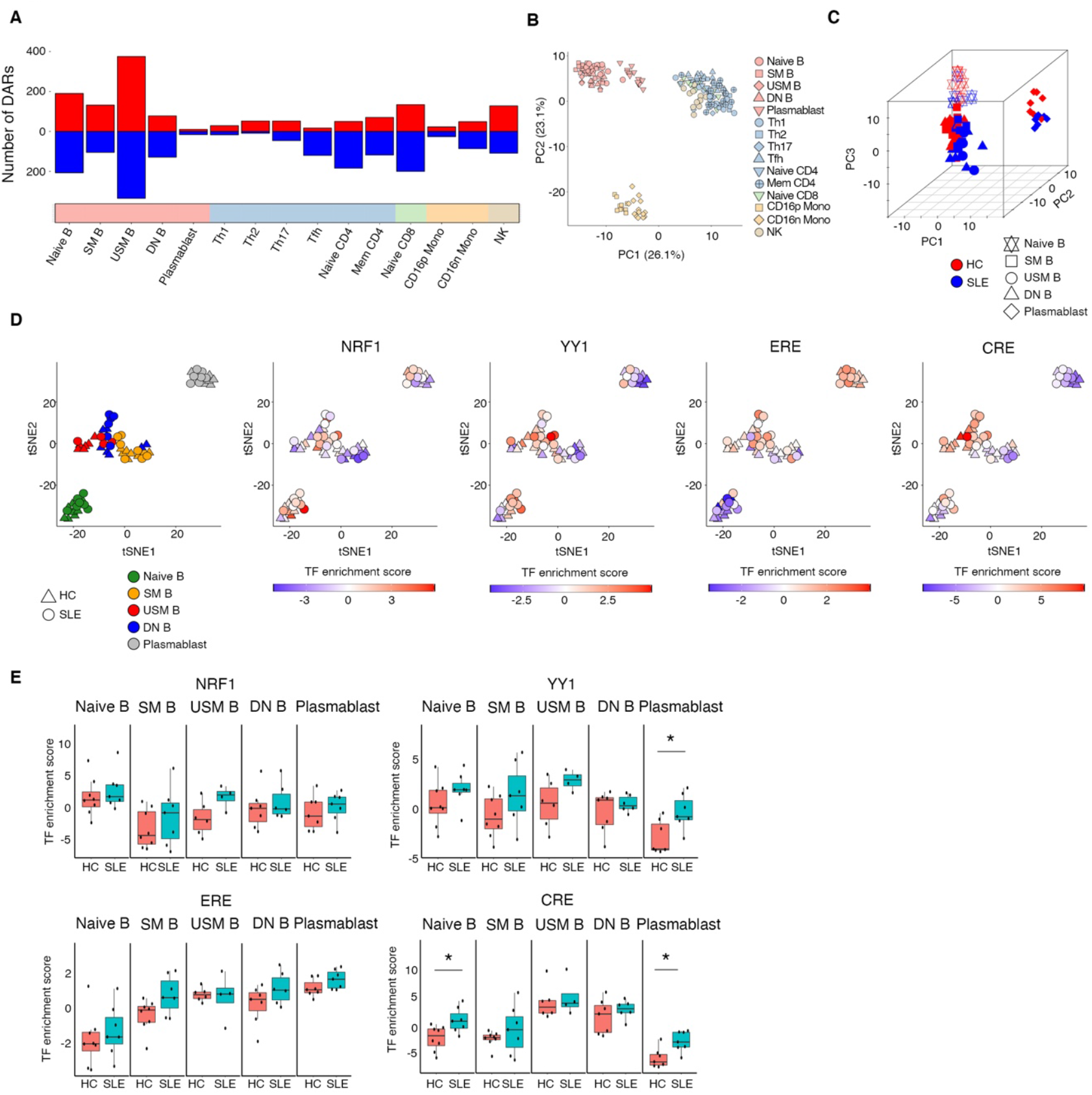
Epigenetic regulation of OXPHOS signature genes according to ATAC-seq analysis. (**A**) Differentially accessible regions were calculated using DiffBind. (**B** and **C**) PCA clustering using all transcription factor enrichment scores calculated by chromVAR. Immune cells were clustered into each parental immune cell subset (B). Data from HCs (red) and SLE patients (blue) were annotated according to each B cell subset (C). (**D**) t-distributed Stochastic Neighbor Embedding (tSNE) clustering of B cell subsets according to the z-score for all the transcription factors (TFs). Each B cell subset was colored (left) by subtypes and each symbol was colored by z-scores for TFs inducing OXPHOS gene expression: NRF1, YY1, ERE, and CRE. HCs are presented as triangles and SLE patients as circles. (**E**) The calculated TF enrichment scores of NRF1, YY1, ERE, and CRE are shown in the box plot. * *p* < 0.05.

### Association between the genetic risk of SLE and an OXPHOS signature by eQTL analysis

GWAS has identified many single nucleotide polymorphisms (SNPs) associated with the risks of several autoimmune diseases. Several approaches have been taken to identify causal genes by integrating risk SNP, eQTL, and transcriptomic data [26,27]. We recently constructed an eQTL database, ImmuNexUT, which consists of 336 subjects with immune-mediated diseases, and 79 HCs (see Methods) [28]. We identified some SLE susceptibility genes with *cis*-eQTL associations (eGenes) from ImmuNexUT by assessing colocalization of GWAS signals and eQTL signals. As this study revealed the importance of OXPHOS in SLE B cells, we picked up mitochondrial function-related 13 genes from these eGenes in B cell subsets (figure 3A and online supplemental table S2). Within these genes, *BANK1, LYST* and *UBE2L3* expression showed strong correlations with the OXPHOS signature in B cell subsets from SLE patients (online supplemental figure S3A). These results suggest the possibility that the OXPHOS signature in B cells co-operates with genetic risk pathway in the pathogenesis of lupus.

**Figure 3.**
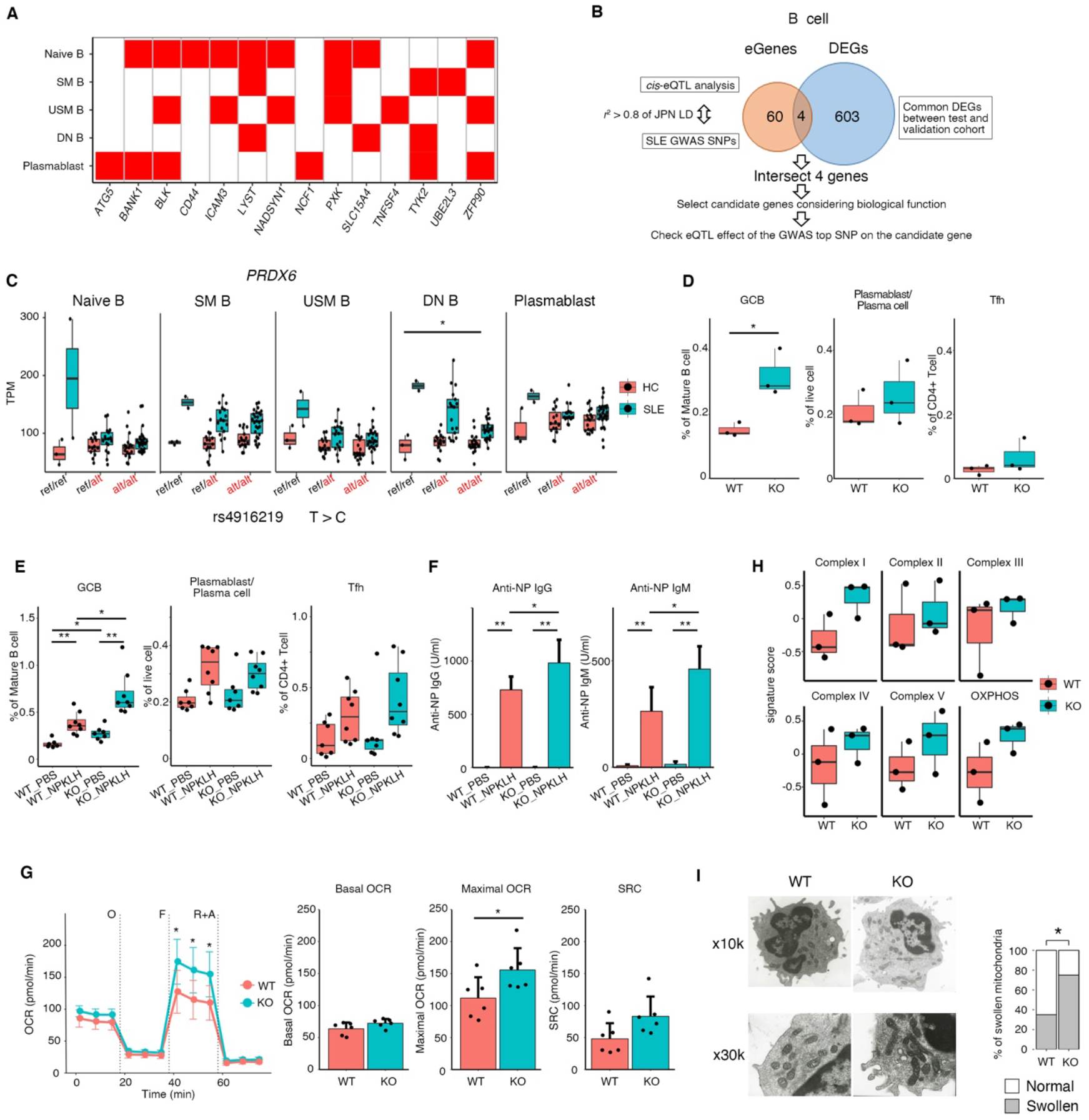
Identification of PRDX6 as a key driver gene in B cells of SLE patients, and analysis of its function using knockout mice. *cis*-eQTL analysis of transcriptomic data from 79 HCs and 336 patients with immune-mediated diseases (IMD) using SLE GWAS Catalog SNP data and Japanese LD information as well as European LD information. (**A**) SLE susceptibility gene list with *cis*-eQTL effects in B cell subsets using our IMD and HC data sets. Red color indicates that a *cis*-eQTL effect exists. (**B**) Schematic diagram of identifying key driver genes in the B cells of SLE patients. (**C**) *cis*-eQTL association analysis of rs4916219 for *PRDX6*. Allele C of rs4916219 of *PRDX6* was identified as a risk haplotype using GWAS data from Japanese SLE patients. ref: reference. alt: alternative. * *p* < 0.05, pink: HCs, turquoise: SLE patients. (**D**) Analyses of GCB, plasmablast, and T follicular helper cell (Tfh) subsets in the spleen of wild type (WT) and *Prdx6* knockout (KO) mice. n = 3. Statistical test was performed using Student’s t-test. * *p* < 0.05. (**E** and **F**) Percentages of GCB, plasmablast, and Tfh cells in the spleen (D) and anti-NP IgG and IgM production (E) in WT and Prdx6 KO mice after NP-KLH immunization. n = 8. (**G**) The basal OCR, maximal mitochondrial respiration rate (maximal OCR) spare respiratory capacity (SRC) and expression of ETC complex and OXPHOS signature genes in the splenic B cells of steady-state *Prdx6* KO mice. n = 6. (**H**) Each complex signature and OXPHOS signature scores were calculated using the splenic B cells of steady-state *Prdx6* KO and WT mice. n = 3. Statistical test was performed using Student’s t-test. * *p* < 0.05. (**I**) Transmission electron microscopy analysis of B cells purified from the spleens of WT and *Prdx6* KO mice. The percentage of cells with swollen (> 500 nm/φ) mitochondria among 20 cells analyzed.

### Peroxiredoxin 6 (*PRDX6*) is a key driver gene in the B cells of SLE patients via regulation of mitochondrial function

Next, we attempted to identify KDGs specific to the B cells of SLE patients referencing the KDG approach [26]. As summarized in figure 3B, we identified SLE-specific eGenes and B cell DEGs between SLE patients and HCs. Of the KDGs overlapping between the eGenes and DEGs (online supplemental table S3), we focused on *PRDX6* as a key gene in B cells, based on its antioxidant functions. As shown in figure 3C and online supplemental figure S4A, the eQTL effect of SLE-risk-associated SNPs led to downregulation of *PRDX6*, suggesting that *PRDX6* expression in B cells has a protective role in SLE pathogenesis [29]. We found a significant increase in the proportion of germinal center B cells (GCBs) in steady-state *Prdx6* knockout (KO) mouse splenocytes (figure 3D and online supplemental figure S4B). Following primary immunization with NP-KLH in alum, the proportion of GCBs was significantly increased in *Prdx6* KO mice compared with wild type (WT) mice, accompanied by upregulated antibody production (figure 3E and F). Splenic B cells lacking *Prdx6* demonstrated an elevated mitochondrial respiration rate (figure 3G), consistent with increased mRNA expression of ETC complex-related and OXPHOS genes in *Prdx6* KO mouse B cells (figure 3H). Mitochondrial disruption was confirmed by the significantly higher proportion of swollen mitochondria in *Prdx6* KO than in WT mouse B cells (figure 3I). These results suggest a protective role of PRDX6 in SLE pathogenesis by negatively regulating plasmablast differentiation via mitochondrial function.

### The OXPHOS gene signature predicts long-term prognosis and is associated with certain clinical phenotypes of SLE

To address long-term risk evaluation in SLE, we examined the associations between the OXPHOS signature and clinical parameters, including the Systemic Lupus International Collaborating Clinics/American College of Rheumatology damage index (SDI), in our cohort. After excluding the patients with chronic kidney disease (see Methods), 74 SLE patients were evaluated in the following analysis. Patients in the test cohort clustered into those with high scores for the OXPHOS signature (see Methods) were significantly enriched among the patients with an SDI > 0 (*p* = 0.03), whereas those with high scores of the type I IFN signaling-related signature showed no enrichment among patients with an SDI > 0 (figure 4A). These enrichments were verified in the validation cohort (*p* = 0.08) (online supplemental figure S5A).

**Figure 4.**
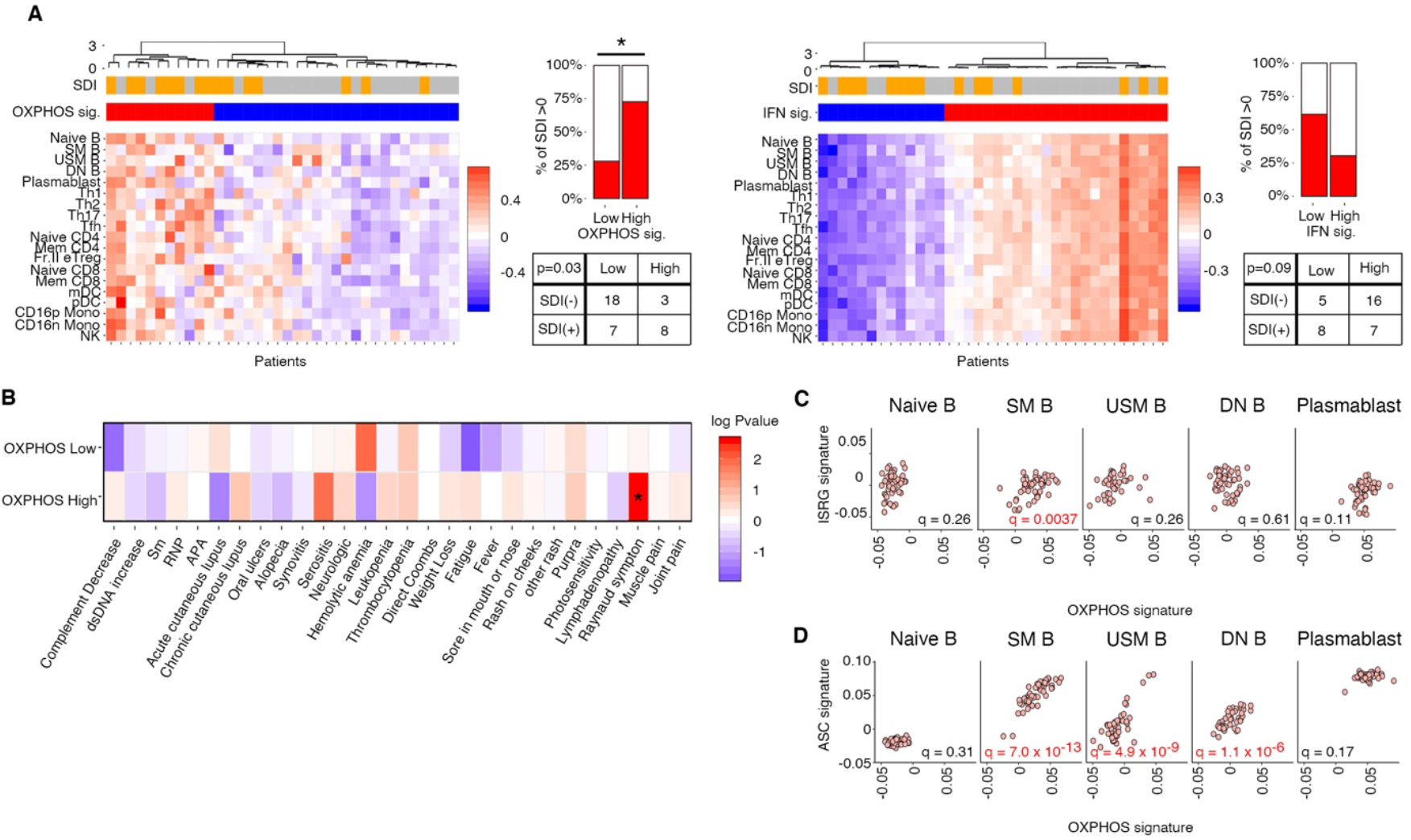
OXPHOS gene signature as a predictor of long-term prognosis in SLE patients in association with clinical phenotypes and its importance in the differentiation of plasmablasts from SLE memory B cells.s. (**A**) Relationships of SDI with the OXPHOS signature score (left) and type I IFN signaling-related signature genes (ISRGs) score (right) in the test cohort (n = 49). SLE patients without chronic kidney disease were clustered according to high- and low-scoring signatures by the hierarchical clustering method. Enrichment analysis of each patient cluster was performed with SDI > 0 and < 0 * *p* < 0.05. (**B**) Clinical characteristics of the patients from the test and validation cohorts in the high/low OXPHOS signature clusters among the patients with SDIs > 0. The data from the test and validation cohorts were combined in this analysis. Fisher’s exact test was used to test for nonrandom associations between two categorical variables, and the signed –log10 *p*-values were visualized by heatmap. * *p* < 0.05. (**C** and **D**) Correlations of the OXPHOS gene signature with type I IFN signaling-related signature (184 genes) (C) and antibody-secreting cell (ASC) (D) gene signatures in SLE B cell subsets.

We investigated which clinical phenotypes correlated with the OXPHOS signature and organ damage. We stratified SLE patients with SDIs > 0 from both cohorts (figure 4B). Patients with Raynaud’s syndrome were significantly enriched in the patients with high OXPHOS signatures and SDIs > 0. Raynaud’s syndrome is a transient and peripheral vasoconstrictive response to cold temperatures. The relationship between the OXPHOS signature and Raynaud’s syndrome might reflect vascular damage in SLE patients.

### Importance of the OXPHOS signature, in association with the type I IFN signaling signature, in plasmablast differentiation in SLE patients

Because the OXPHOS-related module in memory B cells was associated with the IFN signaling module (figure 1D), type I IFN signaling-related genes (ISRGs) may affect the OXPHOS pathway. We evaluated the relationship between the OXPHOS signature score and the ISRGs signature score using 184 genes related to type I IFN signaling (see Methods). The OXPHOS signature score was highest in plasmablasts and was significantly correlated with the ISRGs score in memory B cells, suggesting that the OXPHOS signature is elevated during the differentiation of memory B cells to plasmablasts (figure 4C). These two signatures also showed a correlation in Th1 and memory CD8 T cells, which might reflect increased mitochondrial membrane potential in the T cells of SLE patients (online supplemental figure S5B) [16]. This correlation was confirmed in the validation cohort and was not detected in the HCs (online supplemental figure S5C and D). Notably, the OXPHOS signature score was also significantly correlated with the antibody-secreting cell (ASC) signature score [30] in DN B cells and memory B cells in both cohorts (figure 4D and online supplemental figure S5E). These data indicate that elevated expression of OXPHOS signature genes in the memory B cells of SLE patients is related to type I IFN signaling, leading to plasmablast differentiation.

### A specific type I IFN signature gene set related to OXPHOS, associated with SLE damage accrual

As the OXPHOS signature score showed a correlation with the ISRGs score in memory B cells (figure 4C), we evaluated our transcriptomic data using a hierarchical clustering approach to determine the specific ISRGs that are associated with OXPHOS-related genes in SLE patients. Total six clusters were identified and the reproducibility of the clusters was validated by factor analysis (figure 5A and online supplemental figure S6A) [31]. Among the six gene sets obtained (online supplemental table S4), C6 gene set showed a strong correlation with the OXPHOS signature in both cohorts in all immune cell types (figure 5B, online supplemental figure S6B and C). The correlations of the gene set other than C6 with the OXPHOS signature are presented in online supplemental table S5. SLE patients with a high C6 signature tended to be enriched among those with SDIs > 0 in both test and validation cohorts (*p* = 0.06 and *p* = 0.09, respectively), and joint analysis of two cohorts showed significant enrichment of SLE patients with a high C6 signature among those with SDIs > 0 (*p* = 0.02) (figure 5C). In addition, patients with high expression of C6 signature and SDIs > 0 were characterized by neurologic disorders (figure 5D), indicating that C6 signature predicts a risk of neurologic dysfunction.

**Figure 5.**
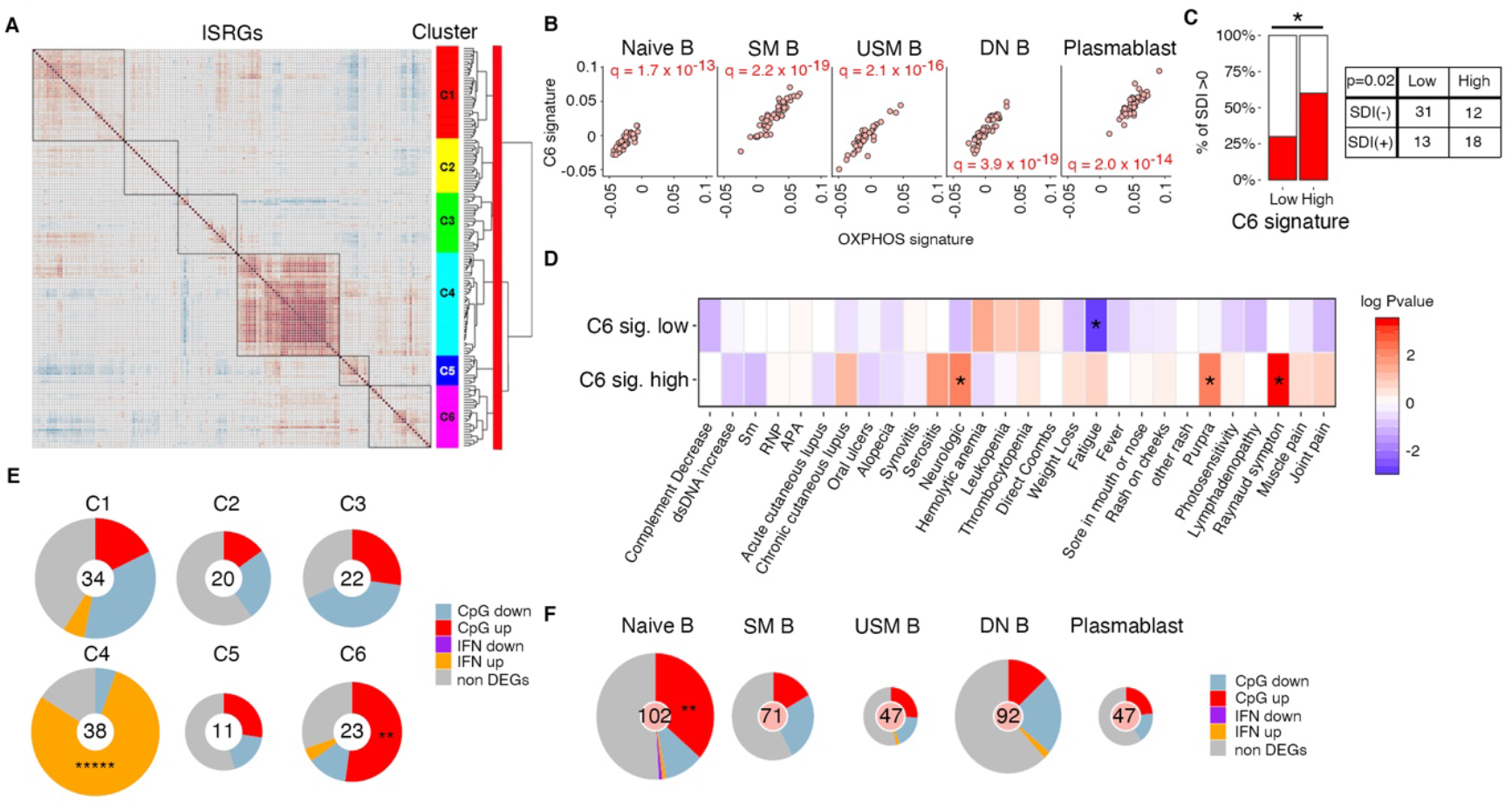
Identifying a specific type I IFN signaling-related gene set with a strong relationship to the OXPHOS gene signature. (**A**) Hierarchical clustering of 184 type I IFN signaling-related genes according to correlation coefficient of their expressions in SLE patients from the test cohort. Six clusters (C1–C6) were identified. (**B**) Correlations between OXPHOS and C6 gene signatures in each B cell subset. (**C**) Relationship between the C6 signature score and SDI in the test and validation cohorts. SLE patients without chronic kidney disease were clustered according to high- and low-scoring signatures by the hierarchical clustering method. Enrichment analysis of each patient cluster by joining both cohorts’ clustering results was performed with SDI > 0 and < 0. * *p* < 0.05. (**D**) The clinical characteristics of the SLE patients from the test and validation cohorts with high/low-scoring C6 signatures and with SDI > 0. Data from the test and validation cohorts were combined in this analysis. Fisher’s exact test was used to test for nonrandom associations between two categorical variables, and the signed –log10 *p*-values were visualized by heatmap. * *p* < 0.05. (**E**) Relationships of each type I IFN signaling pathway gene cluster with DEGs in human memory B cells under TLR9 agonist (CpG) and type I IFN stimulation. Specific DEGs between CpG- or type I IFN-stimulated and un-stimulated human memory B cells were calculated. Each gene cluster was annotated to these DEGs. Red: upregulated DEGs under CpG stimulation; Blue: downregulated DEGs under CpG stimulation; Orange: upregulated DEGs under type I IFN stimulation; Purple: downregulated DEGs under type I IFN stimulation. ** *p* < 0.01, ***** *p* < 0.00001 (**F**) Feature selection using the Boruta package in RNA-seq data from 107 SLE patients. The important features and genes in B cell subsets for distinguishing patients with neurologic disorders were selected. DEGs between CpG- or type I IFN-stimulated and un-stimulated human memory B cells were calculated. Enrichments of each DEGs in these classifier genes were assessed using Fisher’s exact test by comparing the percentages of each DEGs in whole genes analyzed. Each classifier gene was annotated to these DEGs. Red: upregulated DEGs under CpG stimulation; Blue: downregulated DEGs under CpG stimulation; Orange: upregulated DEGs under type I IFN stimulation; Purple: downregulated DEGs under type I IFN stimulation. ** *p* < 0.01

Notably, almost 50% of C6 genes were identified as DEGs in the memory B cells after CpG stimulation versus no stimulation (figure 5E). Most of these DEGs (*GBP2, HMGB1, HSP90AB1, HSPD1, IRF2, POLR2F, POLR2L, POLR3H, POLR3K, UBB, XRCC5*, and *XRCC6*) were upregulated by CpG stimulation. Moreover, feature selection for neurologic disorders using the Boruta algorithm in the expression data from our SLE patients detected a significant enrichment of upregulated DEGs following CpG stimulation in memory B cells within the classifiers in Naive B cells (figure 5F and online supplemental figure S6D). These observations suggest a C6 genes-mediated linkage between innate immune signaling and the progression of neurologic dysfunction. Our results suggest that two gene sets related to TLR-induced signaling, OXPHOS-related genes and C6 genes, are SLE key pathways associated with damage accrual.

## DISCUSSION

We present a precise cell-type-specific multi-omics analyses to identify immunological pathways involved in SLE. Although a previous multi-omics analysis identified several clinically meaningful linkages [32], that analysis was performed in PBMCs or whole-blood samples, representing a combination of immunological modifications from different immune cell subsets. Although recent study on single-cell RNA sequencing of SLE PBMCs has revealed heterogeneity of immune cell subsets precisely [33], our bulk RNA sequencing approach on as many as 18 immune cell subsets had advantages on detecting relatively low-expression genes and differences in gene expressions between each immune cell subset independent of its proportion in PBMCs. Under increasing attention to a treat-to-target approach for SLE [34–36], a critical issue in SLE is identification of immunological pathways related to prognosis.

We identified OXPHOS-related genes as key players in SLE, particularly in memory B cells, the open chromatin status of which demonstrated the highest SLE genetic risk among the immune cell subsets. We revealed a significant association between OXPHOS signature and ASC signature. Our observation is consistent with a previous report showing that upregulation of OXPHOS using dichloroacetate increased proportion of plasmablasts [37]; the importance of mitochondrial reactive oxygen species (mtROS) regulation in plasmablast differentiation has also been reported [38]. The finding that several susceptible genes by a GWAS on SLE were related to mitochondrial function suggests the importance of mitochondrial function in SLE pathogenesis [39]. In addition, we identified PRDX6 as a key driver to cellular metabolism in the B cells of SLE patients. PRDXs represent a superfamily of non-selenium peroxidases that catalyze the reduction of peroxides. Although previous studies on the function of PRDX6 in autoimmune mouse models reported controversial findings [40–42], our *cis*-eQTL analysis and *Prdx6* KO mice indicated that PRDX6 in B cells protects against SLE. The effect size of PRDX6 on SLE pathogenesis via regulation of B cells remains to be clarified; however, the combined effect of PRDX6 impairment and activation of the innate immune system may lead to the SLE phenotype.

Regarding the clinical aspects of SLE, the OXPHOS signature was related to SDI, an indicator of damage accrual in our SLE cohorts. Oxidative stress is widely accepted as a biomarker of disease activity and organ damage in various pathologies, such as cardiovascular disease. Panousis *et al*. recently reported that genes related to OXPHOS were enriched among DEGs between SLE patients and HCs in PBMCs, and were closely associated with activity and severity in SLE [3]. We also found the OXPHOS signature association with Raynaud’s syndrome, which was related to mtROS in vascular smooth muscle cells [43], and focal involvement of the central nervous system in SLE [44,45]. Our result suggested that the OXPHOS signature is associated with not only activity of SLE, but also susceptibility of SLE supported by genetic risk. Because the OXPHOS signature in B cells was induced by TLR signaling, not type I IFN, persisting innate immune signaling even in low disease activity may influence the long-term prognosis of SLE.

We also identified specific TLR signaling related genes that strongly correlate with the OXPHOS signature gene set. In terms of organ damage, a high score of these signature genes was associated with SDI and neurologic disorders, again suggesting linkage between innate immune signaling and organ damage. Our observation also supports crosstalk between TLR-mediated innate immune and inflammasome signaling pathways in the pathogenesis of neuroinflammation [46].

Several limitations of our study should be considered. The study cohort included only Asian patients who had nearly stable disease and were treated with low-dose corticosteroids. We excluded patients under high dosage of steroids because of potentially strong effects on the transcriptome, which might obscure the pathogenic changes in immune cell subsets. Therefore, our analysis may focus on a ‘susceptibility’ signature that persists in the presence of limited disease activity under treatment [3].

Our multi-omics approach in each immune cell type revealed the importance of OXPHOS in memory B cells for SLE progression. We suggest the clinical significance of the OXPHOS signature, which was strongly associated with innate immune signaling and damage accrual in SLE patients. We propose that innate immune signaling including OXPHOS signature genes, as new treatment targets and long-term prognostic markers of SLE.

## MATERIALS AND METHODS

See online supplementary materials and methods.

## Acknowledgments

The super-computing resource was provided by Human Genome Center, Institute of Medical Sciences, The University of Tokyo (http://sc.hgc.jp/shirokane.html). We thank Mr. Satoru Fukuda (Department of Pathology, Graduate School of Medicine, University of Tokyo) for his technical assistance with the transmission electron microscopy analysis. We also thank the following people for technological expertise and support: Ms. J. Takezawa and Ms. K. Watada.

## Funding

This study was supported by Chugai Pharmaceutical Co., Ltd., Tokyo, Japan, the Center of Innovation Program from Japan Science and Technology Agency (JST) (JPMJCE1304), the Ministry of Education, Culture, Sports, and the Japan Agency for Medical Research and Development (AMED) (JP17ek0109103h0003).

## Author contributions

Y.T. performed and analyzed the majority of experiments in this study. Y.I. designed the experiments, conducted analyses, as well as wrote the manuscript. M.N. provided in vitro support. Y.N. made suggestions on ATAC-seq analysis. M.O. performed eQTL analysis. S.S., T.O., K.I., A.S., and Y.K. provided technical assistance. K.Y. and K. F. supervised the project and co-wrote the manuscript.

## Competing interests

Y.T., M.O., Y.N. and T.O. belongs to the Social Cooperation 866 Program, Department of functional genomics and immunological diseases, supported by 867 Chugai Pharmaceutical. Y.N. is an employee of Chugai 865 Pharmaceutical. K.F. receives consulting honoraria and research support from 868 Chugai Pharmaceutical.

## Data and materials availability

All analyzed sequencing datasets and open chromatin data, during the current study were deposited in the National Bioscience Database Center (NBDC) Human Database (http://humandbs.biosciencedbc.jp/) with the accession code hum0214, which can be downloaded upon request. We used publicly available software for the analyses.

## Key messages

### What is already known about this subject?

- Systemic lupus erythematosus (SLE) is an autoimmune disease that can affect various organs. Both adaptive and innate immune system contribute to SLE pathogenesis, but precise mechanism of immune system regulation remains unclear. The type I interferon (IFN) signature is a prominent feature of SLE, but its correlation with SLE activity is controversial and it cannot predict disease prognosis.

### What does this study add?

- Our integrative transcriptomic, epigenetic, and genomic analysis on 18 immune cell subsets from PBMC of SLE patients revealed the importance of memory B cells via oxidative phosphorylation (OXPHOS)/mitochondrial dysfunction.
- By combining differentially expressed genes (DEGs) analysis and expression quantitative trait loci (eQTL) analysis, *PRDX6*, one of the SLE susceptibility genes, was picked up as a candidate key driver gene for SLE pathogenesis. Mitochondrial respiration in B cells as well as antibody production were upregulated by *Prdx6* deficiency.
- The OXPHOS signature in patients with SLE could predict long-term prognosis and was associated with certain clinical phenotypes. Additionally, we revealed that the gene set related to toll-like receptor signaling was strongly correlated with OXPHOS signature, suggesting innate immune signal importance for SLE pathogenesis.

### How might this impact on clinical practice or future developments?

- Our immune cell multi-omics analysis proposed OXPHOS signature along with innate immune signaling as a new treatment target in SLE B cells. Furthermore, OXPHOS signature could be a long-term prognostic marker of patients with SLE.

---

**Online Supplemental Figure S1.**
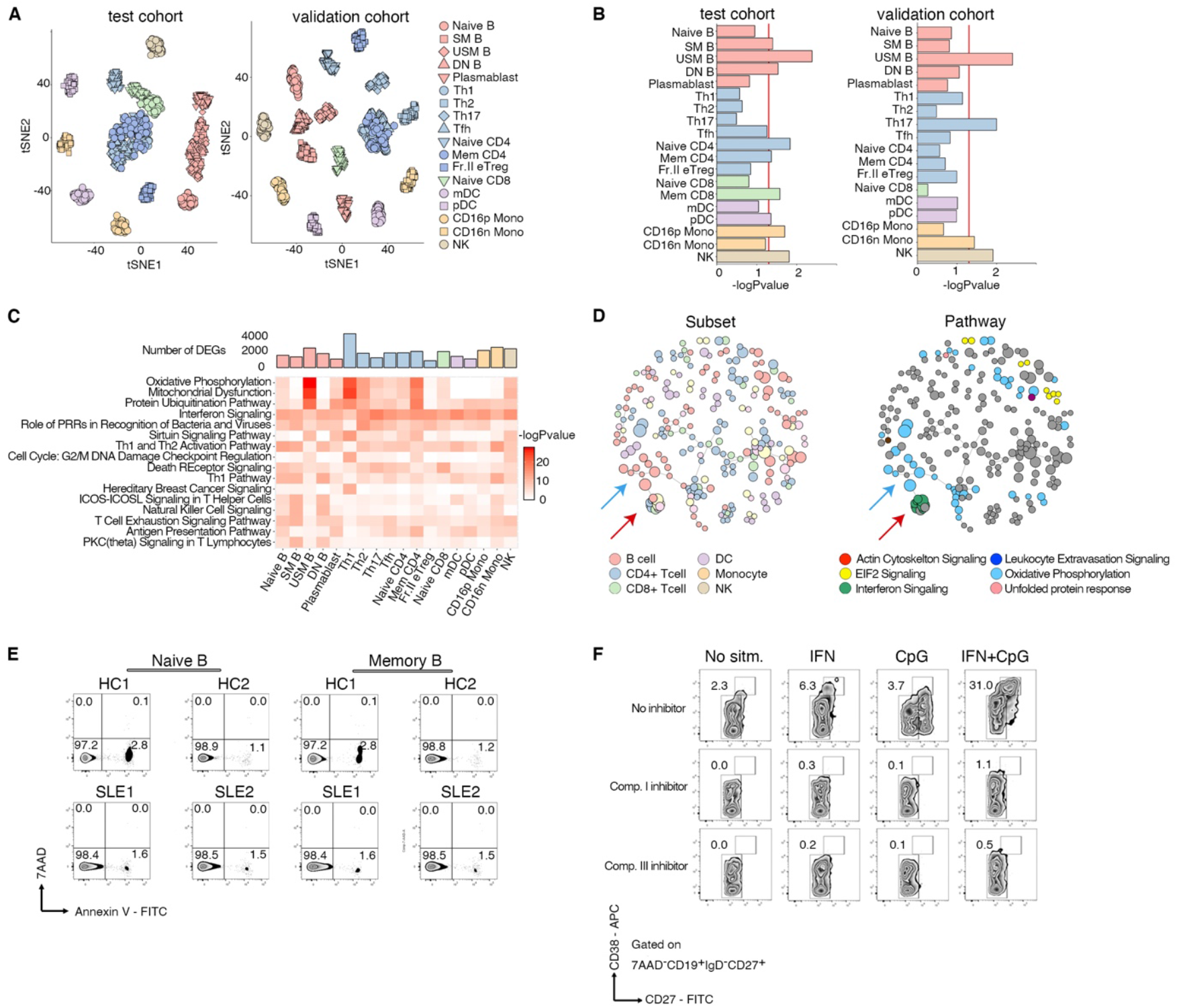
Quality control of our transcriptomic data and the importance of memory B cells in SLE pathogenesis. (**A**) Data quality control and removal of outliers. Gene expression was normalized using the TMM method. The samples with an average *r*^2^ < 0.8 were omitted as outliers. The remaining samples were clustered by t-distributed stochastic neighbor embedding (tSNE). (**B**) Linkage disequilibrium score regression analysis of the top 1,000 DEGs using summary statistics from a GWAS of SLE in each immune cell subset in both the test (left) and validation (right) cohorts. Red line indicates significance at *p* < 0.05. (**C**) Pathway analysis of DEGs in all immune cell subsets with *q* < 0.001 in the validation cohort. The pathways with a –log10 *p*-value > 10 in any immune cell subset were visualized by heatmap. (**D**) Network analysis of each module correlation, as in Fig. 1D, in the validation cohort. Only the modules with *r*^2^ > 0.8 were visualized. Red arrow indicates IFN signaling modules. Blue arrow indicates OXPHOS modules in memory B cells. (**E**) Human naïve and memory B cells magnetically sorted from PBMCs of two SLE patients and two HCs using MojoSort™ (BioLegend). Cells were stained with both Annexin V and 7-AAD, and the percentage of pro-apoptotic Annexin V-positive and 7-AAD-negative cells was assessed. (**F**) Inhibitory effects of ETC genes on plasmablast differentiation. Human memory B cells were purified from PBMCs and cultured for 72 h with combinations of CpG ODN 2006 and IFN-α, CpG ODN 2006 (2.5 mg/ml) and recombinant human IFN-α (1,000 U/ml) were used. The following ETC inhibitors were used: 5 mM rotenone for complex I and 1 mM antimycin A for complex III.

**Online Supplemental Figure S2.**
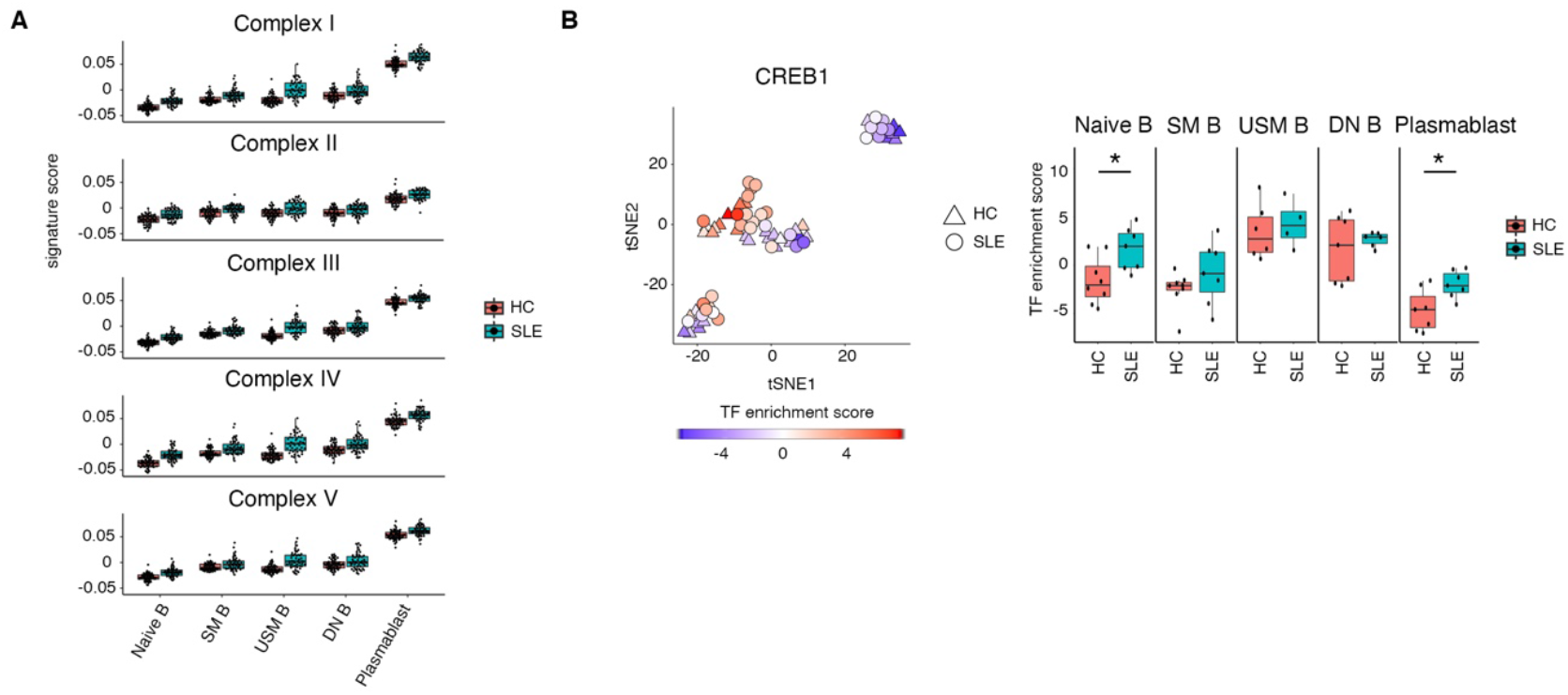
Differentially accessible regions of each immune cell subset between SLE patients and HCs and enrichment analysis of transcription factors regulating ETC genes expression. (**A**) mRNA expression of each ETC gene in SLE patients versus HCs. Each ETC gene signature score was significantly (*p* < 0.05) elevated in SLE patients. The genes used to calculate expression are listed in Table S1. (**B**) tSNE clustering of B cell subsets according to the transcription factor enrichment score of CREB1. Each B cell subset is colored by subtypes and each symbol is colored by z-scores for CREB1. The enrichment scores of each B cell subset are also presented in the box plot. HCs are presented as triangles and SLE patients as circles.

**Online Supplemental Figure S3.**
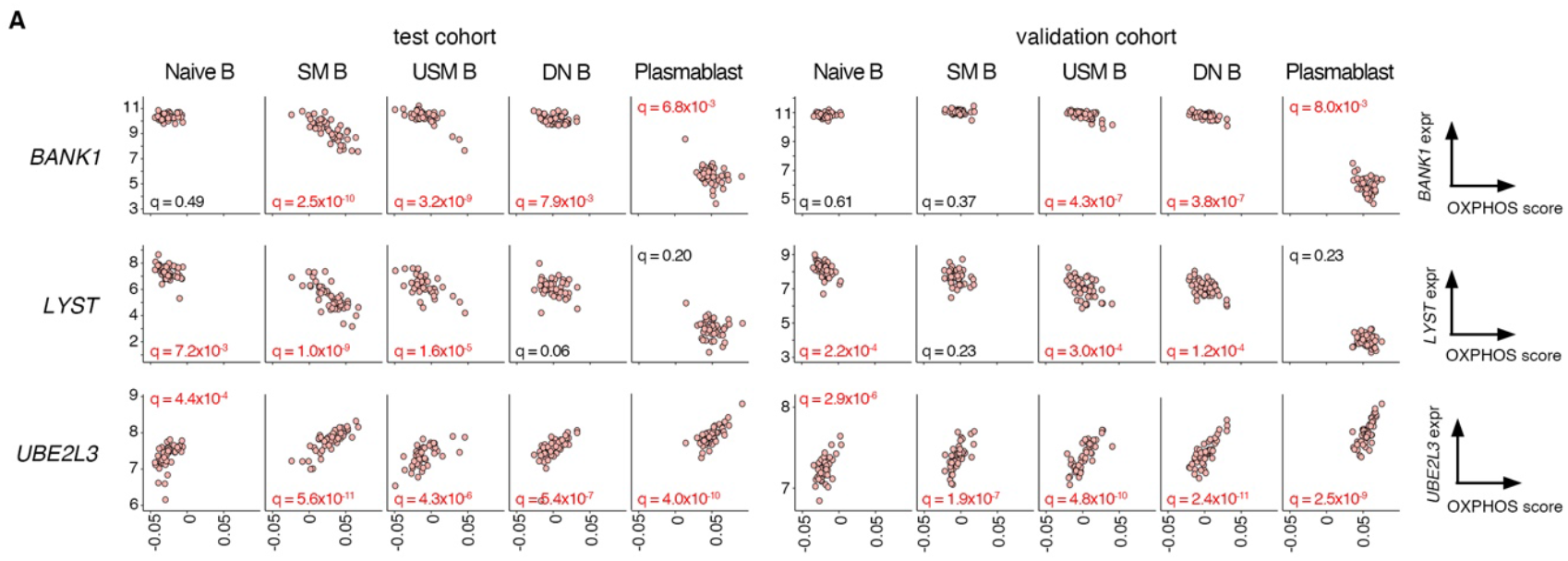
Genetic regulation of OXPHOS signature genes according to *cis*-eQTL analysis. (**A**) Correlations of the OXPHOS signature with the expression levels of *BANK1, LYST* and *UBE2L3* in B cell subsets. Test cohort (left) and validation cohort (right). *q*-values under 0.05 were shown in red.

**Online Supplemental Figure S4.**
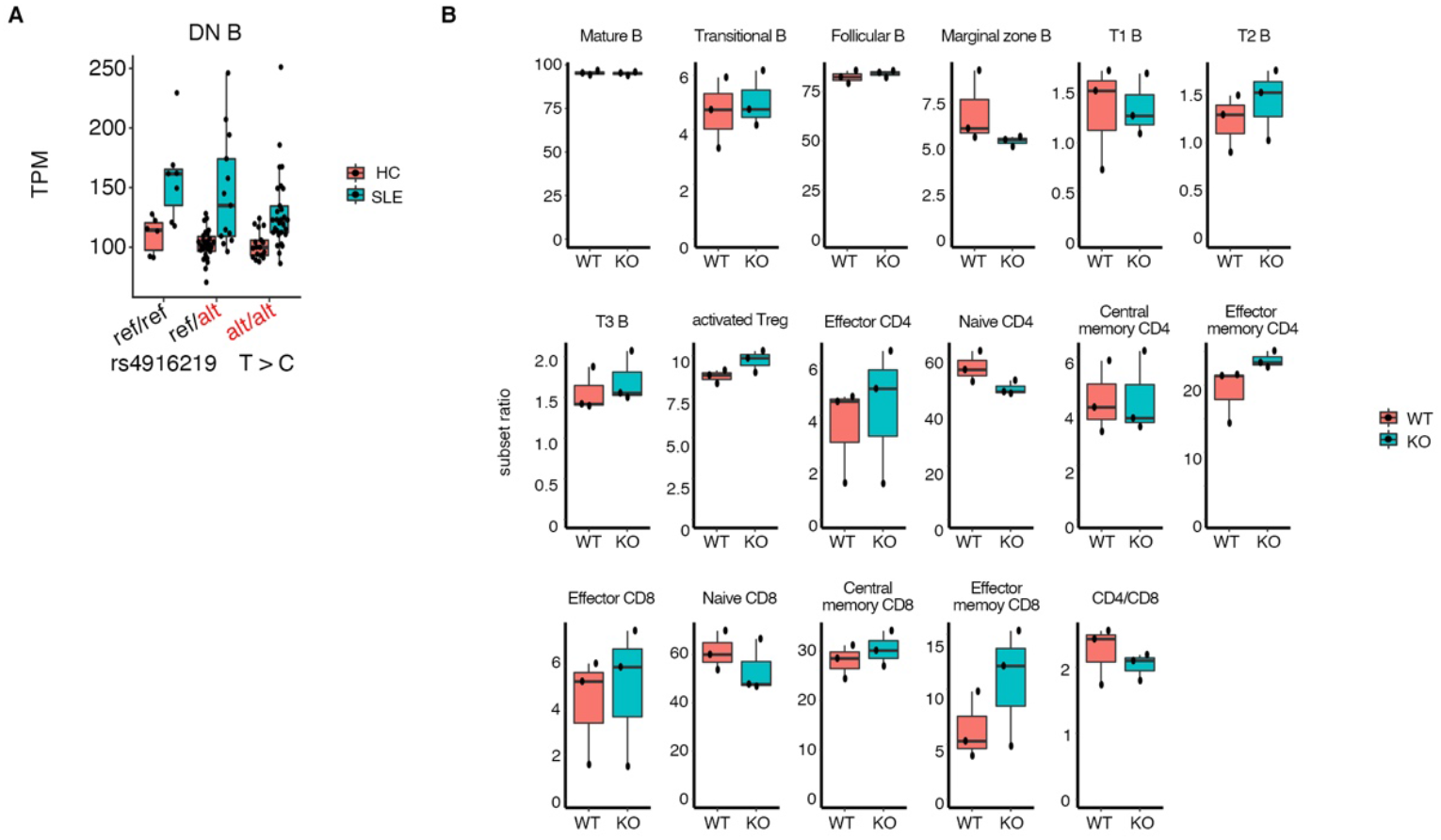
The *cis*-eQTL effect of PRDX6 on B cells in our validation cohort and analysis of splenic immune cell subsets in Prdx6 KO mice. **(A)** *cis*-eQTL association analysis of rs4916219 for *PRDX6* in DNB of our validation cohort. Allele C of rs4916219 of *PRDX6* was identified as a risk haplotype using GWAS data from Asian SLE patients. ref: reference. alt: alternative. (**B**) Proportion of each immune cell subset to each parent subset was calculated in the spleens of *Prdx6* KO mice. n = 3. Statistical tests were performed using Student’s t-test.

**Online Supplemental Figure S5.**
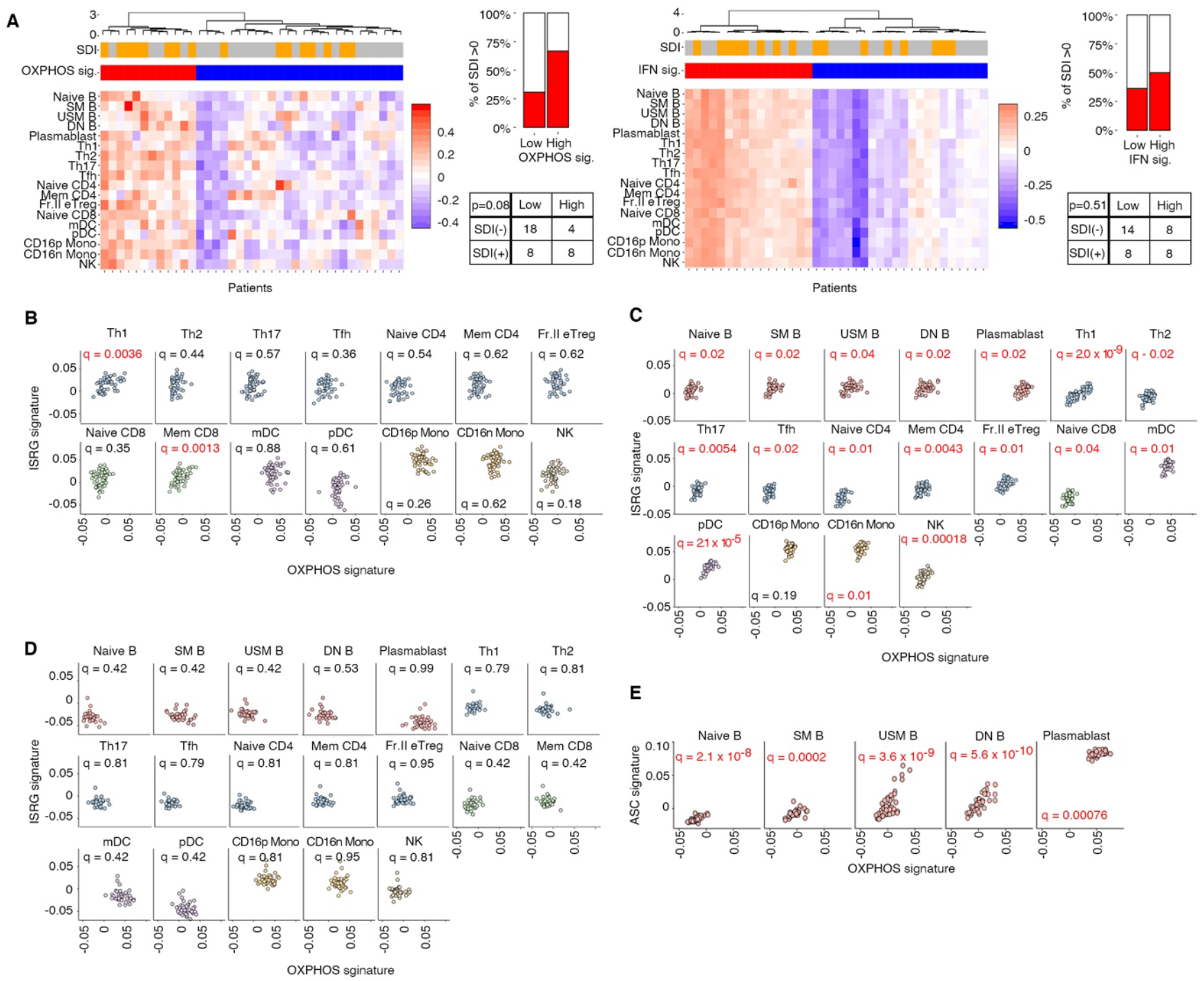
Reproducibility of the relationships of the OXPHOS and type I IFN signaling-related signature scores with SDI in the validation cohort, and relationships of the OXPHOS signature score with the type I IFN signaling-related and ASC signature scores. (**A**) SLE patients from the validation cohort were clustered according to the score of the OXPHOS or type I IFN signaling-related genes (ISRGs) signature. Enrichment analysis of each patient cluster was performed according to SDI > 0 versus < 0. (**B**, **C**, and **D**) Correlation between the OXPHOS and ISRGs signature scores in immune cell subsets other than B cells (B), in the validation cohort (C), and in HCs (D). (**E**) Correlation between the OXPHOS and ASC signature scores in B cell subsets in the validation cohort. *q*-values under 0.05 were shown in red.

**Online Supplemental Figure S6.**
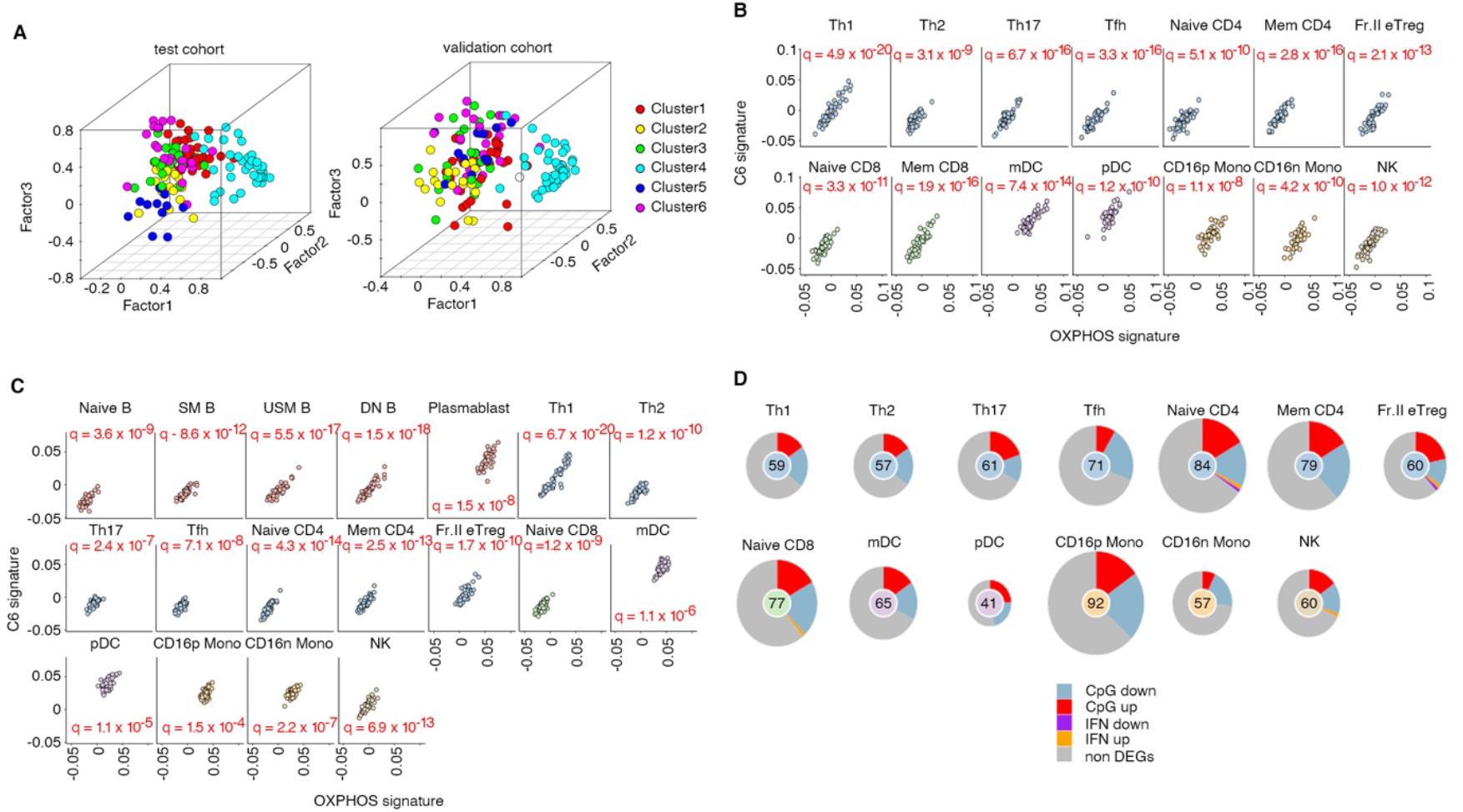
The OXPHOS signature was related to a specific type I IFN signaling-related gene set under the control of innate immune signaling. **(A)** Factor analysis of the expression of multiple type I IFN signaling-related genes in SLE patients. C1–6 genes identified in Fig. 6A are circled in each color. Red: Cluster1; Yellow: Cluster2; Green: Cluster3; Light blue: Cluster4; Blue: Cluster5; Purple: Cluster6. **(B)** Correlation between OXPHOS and C6 signatures in each immune cell subset other than B cells in the test cohort. **(C)** Correlation between OXPHOS and C6 signatures in all immune cell subsets in the validation cohort. **(D)** Feature selection using the Boruta package in RNA-seq data from 107 SLE patients. The important features and genes in immune cell subsets other than B cells for distinguishing patients with neurologic disorders were selected. DEGs between CpG- or type I IFN-stimulated and un-stimulated human memory B cells are shown. Enrichments of each DEG in these classifier genes were assessed using Fisher’s exact test by comparing the percentages of each DEG to whole genes analyzed. Each classifier gene is annotated to these DEGs. Red: upregulated DEGs under CpG stimulation; Blue: downregulated DEGs under CpG stimulation; Orange: upregulated DEGs under type I IFN stimulation; Purple: downregulated DEGs under type I IFN stimulation. *q*-values under 0.05 were shown in red.

## Notes

### Competing Interest Statement

Y.T., M.O., Y.N. and T.O. belong to the Social Cooperation Program, Department of functional genomics and immunological diseases, supported by Chugai Pharmaceutical. Y.N. is an employee of Chugai Pharmaceutical. K.F. receives consulting honoraria and research support from Chugai Pharmaceutical.

