## Supplementary information for "Immune cell multi-omics analysis reveals contribution of oxidative phosphorylation to B cell functions and organ damage of lupus"

**Online supplementary materials and methods**

**Study design**

The purpose of this study on SLE was to unveil the contribution of each immune cell subset to its pathogenesis by investigating genetic, epigenetic, and transcriptional regulation on immune cells. In addition, we aimed to find clinical meanings as well as importance of the results. To this end, we generated and analyzed RNA-seq, ATAC-seq, and genomic data from human peripheral blood samples. SLE patients with stable activities were recruited in order to avoid strong interferon and/or inflammatory signal overwhelming some lupus susceptibility signatures.

**Human Participants**

In this study, 49 SLE patients and 37 age- and sex-matched HCs for the test cohort and 58 SLE patients and 55 HCs for the validation cohort were recruited. The SLE patients were ambulatory patients at the University of Tokyo Hospital. Ethical approval of this study was obtained from the Human Genome, Gene Analysis Research Ethics Committee of the Graduate School of Medicine and Faculty of Medicine, University of Tokyo (reference number G10095), and informed consent was obtained from all subjects. Eligible patients were > 18 years of age and had been diagnosed with SLE according to the American College of Rheumatology 1997 revised classification criteria. The inclusion criteria were as follows: (1) treatment with < 10 mg/day prednisolone, (2) absence of autoimmune diseases other than Sjögren's syndrome, and (3) absence of malignancies within the last 5 years. All of the patients and HCs were East Asian. To evaluate the effect of the OXPHOS gene signature on long-term prognosis, patients with an estimated glomerular filtration rate < 50 ml/min were excluded, resulting in 36 and 38 SLE patients remaining in the test and validation cohorts, respectively. The clinical characteristics of the SLE patients were assessed with the 2012 Systemic Lupus International Collaborating Clinics criteria [1] and systemic lupus activity measure-revised [2], and are summarized in online supplementary table S6.

**Animals**

C57BL6/J mice were purchased from Japan SLC (Hamamatsu), and *Prdx6* KO mice were cryo-recovered from embryos in the KOMP (KNOCKOUT MOUSE PROJECT) repository. All mice were used at 7–10 weeks of age. All animal experiments were conducted in accordance with institutional and national guidelines.

**Separation of immune cell subsets by multi-parameter flow cytometry**

Human PBMCs were isolated from 25 ml whole blood by centrifugation (1,000 g for 10 min.) over Ficoll, and each immune cell subset was sorted using a flow cytometer (MoFlo XDP [Beckman Coulter] for the test cohort and FACSAria Fusion [BD Biosciences] for the validation cohort). For the definitions of each immune cell subset, we followed the strategy recommended by the Human Immunology Project Consortium [3]. We collected 5,000 cells per subset. The precise gating strategy and list of antibodies used are presented in Table S7.

**RNA-seq library preparation and sequencing**

For the test cohort, total RNA was isolated from each immune cell subset in 350 μl RLT Buffer (Qiagen) containing 1% β-mercaptoethanol. For the validation cohort, total RNA was isolated from each immune cell subset using the MagMAX kit (Thermo Fisher Scientific) following the manufacturer's protocol. RNA quality was assessed by the RNA integrity number obtained from the Agilent BioAnalyzer 2100. Sequencing libraries were constructed from total RNA using Smart-Seq v2 [4] for the test cohort and the Smart-Seq v4 Ultra Low Input RNA Kit for Sequencing (Clontech) for the validation cohort. Paired-end sequencing was performed on the HiSeq 2500 sequencer (Illumina) with a target of approximately 10 million read pairs per sample. The resulting bcl files were deconvoluted and converted to FASTQ format using bcl2fastq v1.8.4 from Illumina. FASTQ files were aligned to the human genome within the UCSC Genome Browser (GRCh38; GenBank assembly GCA_000001405.18) using STAR (v2.5). HTSeq-count (v0.6.1) was used to generate gene counts. In the quality-control analysis, samples with a Phred quality score > 20 were selected using the FASTX-Toolkit (v0.0.14). As the level of mitochondrial transcription is an indicator of cell stress, we applied a cutoff percentage of mitochondrial gene transcripts of < 8% [5]. Genes were filtered to include those with raw read counts ≥ 10 in at least 10% of each cell subset library. The final data set included 18 immune cell subsets composed of approximately 12,000 genes. Genes were normalized using the trimmed mean of M value (TMM) method. For detecting outlier samples, Spearman's correlation for each subset was calculated, and samples with an average $r^2 < 0.8$ were omitted as outliers.

**RNA-seq in stimulated mouse splenic and human B cells**

Using Smart-Seq2, RNA-seq libraries were constructed from mouse splenic B cells purified from WT or *Prdx6* KO mice and human naive/memory B cells stimulated with type I IFN and/or a TLR9 agonist. Paired-end sequencing was performed using the MiSeq sequencer (Illumina) with a target of approximately 300,000 reads per sample. The resulting bcl files were deconvoluted and converted to FASTQ format using bcl2fastq v1.8.4 from Illumina. FASTQ files were aligned to the human genome (GRCh38; GenBank assembly GCA_000001405.18) or mouse genome (GRCm38; GenBank assembly GCA 000001635.2) within the UCSC Genome Browser using STAR (v2.5.3a).

**Fast-ATAC-seq**

An additional eight SLE patients and eight HCs were enrolled for the ATAC-seq analysis. The inclusion criteria for these subjects were the same as those for the transcriptome analysis. We used a Fast-ATAC-seq protocol optimized for blood cells [6]. Five thousand sorted cells in wash buffer (PBS containing 1% FCS and 1 mM EDTA) were pelleted by centrifugation at 500 × g for 5 min at 4°C in a precooled fixed-angle centrifuge. All of the supernatant was removed. Next, transposase mixture (25 μl 2× TD buffer, 2.5 μl TDE1, 0.25 μl 2% digitonin, and 22.25 μl nuclease-free water) (Nextera DNA Library Prep Kit, Illumina; G9441, Promega) was added to the cells, and the pellet was disrupted by pipetting. The transposition reactions were incubated at 37°C for 30 min in an Eppendorf ThermoMixer with agitation at 300 rpm. The resulting DNA was purified using the QIAGEN MinElute Reaction Cleanup kit (28204), and purified DNA was eluted in 10 μl elution buffer (10 mM Tris-HCl, pH 8). Transposed fragments were amplified and purified as described previously[7,8]. Libraries were quantified using qPCR prior to sequencing. All Fast-ATAC-seq libraries were sequenced using paired-end, single-index sequencing on the HiSeq 2500. Dendritic cell subsets (myeloid and plasmacytoid dendritic cells) were not collected for the ATAC-seq analysis because of the limited volume of total blood collected from each subject. We evaluated a total of 15 types of immune cells.

**ATAC-seq data analysis**

Reads were trimmed using a custom script and aligned using Bowtie 2 (v2.3.4.3). Quality control was performed using ATACseqQC (v1.2.10) [9]. For peak calling, data were grouped according to each unique cell type, and peak summits were called using MACS2 (v1.4.3) and filtered using a custom blacklist, as described previously [10]. The complete data set comprised a total of 278,973 peaks. To obtain normalized fragment counts, which were used for all downstream processing steps, coverage tracks were visualized using the integrative genomics viewer [11]. We performed stratified LDSC (available as open-source software at http://www.github.com/bulik/ldsc) to determine whether heritability is enriched in the open chromatin regions in each immune cell subset, using our ATAC-seq peaks for regression and summary statistics from a previous SLE GWAS [12] for the reference panel. We used DiffBind (v2.4.8) (Stark R, Brown G (2011). *DiffBind: differential binding analysis of ChIP-Seq peak data*) to obtain DARs and HOMER (v4.11.1) to identify the genes nearest to the DARs. ChromVAR, which is available as open-source software (http://www.github.com/GreenleafLab/chromVar), was used to compute bias-corrected accessibility and the z-score for each TF annotation in each immune cell subset [13]. We utilized downstream applications for cell clustering using bias-corrected z-scores.

**DEG analysis**

Genes with low read counts were removed before the DEG analysis. DEG analysis was performed using the data sets for each immune cell subset from SLE patients and HCs. We used the edgeR (v3.18.1) package to calculate DEGs; run batch effects were removed, and raw count data were normalized by TMM. Resulting *p*-values were adjusted for multiple hypothesis testing and filtered to retain DEGs with a *q*-value < 0.05. We performed pathway analysis of the DEGs using Ingenuity Pathway Analysis, and the pathways with a −log10 *p*-value > 10 in any immune cell subset were visualized by a heatmap. Heritability estimates of the top 1000 DEGs in each immune cell subset were calculated using LDSC analysis of specifically expressed genes [14]. Genome locations of the DEGs were identified using PBMC data (E062) from the ROADMAP project after excluding quiescent sites. We used summary statistics from a previous GWAS on SLE [15] for our reference panel.

**Module definition using WGCNA**

WGCNA (v1.67) is an open-source package for R available at https://horvath.genetics.ucla.edu/html/CoexpressionNetwork/Rpackages/WGCNA/. We removed genes with a read count < 10 in > 90% of the samples and normalized the data using TMM. Read counts were transformed to logCPM values, and batch effects in the RNA extraction method (due to different columns used for RNA extraction) were removed by an empirical Bayesian framework (comBat in the sva package, v3.24.4). We performed singular value decomposition (SVD) and calculated module eigengenes, defined as the first left singular vector of the expression matrix corresponding to the module. Correlations of each vector value in each module were calculated, and we visualized all correlations of $r^2 > 0.7$ or 0.8 using igraph (v1.2.4) in the R package. The node size represents the relative number of genes in the module, and the presence of an edge indicates a significant correlation between two modules, with the edge thickness reflecting the strength of the correlation. To calculate the signature scores for the gene sets, we also applied SVD and defined the vector value as the score. The score for the OXPHOS gene signature was calculated using the HALLMARK OXIDATIVE PHOSPHORYLATION gene set (200 genes), and that for the type I IFN signaling-related gene (ISRG) signature was calculated based on GO gene sets from MSigDB (v6.1) (response to type I IFN (GO:0034340), regulation of type I IFN production (GO:0032479), and regulation of type I IFN-mediated signaling pathway (GO:0060338): 184 genes, but we excluded those with low expression, leaving 148 genes.) Signature scores were calculated using transcriptomic data from SLE patients.

**Imputation and genotyping**

In the test cohort, genotyping was performed using the Infinium® OmniExpressExome-8 v1.4 BeadChip (GRCh37). For quality control, identity-by-descent estimation of relatedness is used to help phase the individuals accurately over long stretches. In addition, we checked sex and removed monomorphic alleles or alleles with a missing rate > 0.01. After removing outliers by principal component analysis using HapMap populations, we performed pre-phasing-based imputation using the programs SHAPEIT and IMPUTE2 together. To fully account for the uncertainty in imputed genotypes, an observed data likelihood approach implemented in SNPTEST was used, in which the contribution of each possible genotype is weighted by its imputation probability. We utilized liftOver (available as open-source software at http://genome.ucsc.edu/cgi-bin/hgLiftOver) to convert GRCh37 to GRCh38. In the validation cohort, whole genome was sequenced on Illumina's HiSeq X. Data processing was performed based on the standardized best-practice method proposed by GATK (v 4.0.6.0). A total of

6,165,701 variants (minor allele frequency ≥ 5%) in the test cohort and 8,105,611 variants (minor allele frequency ≥ 1%) in the validation cohort were selected and used for eQTL analysis.

#### *cis*-eQTL analysis in HCs and patients with immune-mediated diseases (IMD)

In this analysis, we used our RNA-seq and whole genome sequencing data from 79 HCs, 25 patients with Anti-Neutrophil Cytoplasmic Antibodies (ANCA)-associated vasculitis, 18 patients with adult-onset Still's disease, 23 patients with Behcet's disease, 19 patients with mixed connective tissue disease, 65 patients with inflammatory myositis, 24 patients with rheumatoid arthritis, 18 patients with Sjögren's syndrome, 62 patients with SLE, 67 patients with systemic sclerosis, and 16 patients with Takayasu arteritis [16]. We excluded data from third-degree siblings. After sample QC, low-expressed genes (< 5 count in more than 80% samples or < 0.5 CPM in more than 80% samples) were filtered and residual expression data were normalized between samples with TMM, converted to counts per million, and then normalized across samples using an inverse normal transform. Probabilistic Estimation of Expression Residuals (PEER) method was applied to normalized expression data to infer hidden covariates. Top two genetic principal components, sorter batch, disease types, sex and hidden covariates estimated by PEER were utilized as covariates for eQTL analysis. eQTL analysis was performed using the QTLtools package. This *cis*-eQTL analysis revealed an average of 6,000–8,000 SNP–gene associations in each immune cell subset with a < 0.05 false discovery rate. We used lead SLE-related SNPs from the GWAS Catalog (https://www.ebi.ac.uk/gwas/) and applied two LD data: Japanese LD data and European LD data from the 1000 Genomes Project phase 3. We selected eGenes with $r^2 > 0.8$ at both LD data. A manual document retrieval system was applied to eGenes in any B cell subset using PubMed data base to search their functions in B cells.

### Key driver gene analysis

For disease-related *cis*-eQTL analysis, we used our RNA-seq and genotyping data from 48 SLE patients in test cohort. We excluded data from third-degree siblings. After sample QC, low-expressed genes (< 10 count in more than 90% samples or < 0.1 FPKM in more than 80% samples) were filtered and residual expression data were normalized between samples with TMM, converted to counts per million, and then normalized across samples using an inverse normal transform. PEER method was applied to normalized expression data to infer hidden covariates. RNA extraction batch, sex, age and hidden covariates estimated by PEER were utilized as covariates for eQTL analysis. eQTL analysis was performed using the QTLtools packages (version 1.2). We selected lead SLE-related SNPs from the GWAS Catalog (https://www.ebi.ac.uk/gwas/) and INSIDEGEN (http://insidegen.com/insidegen-home.html). We identified eSNP-eGene lists focusing on the GWAS top SNPs that have *cis*-eQTL effects in strong Japanese LD ($r^2 > 0.8$) within 100kb from TSS sites. Then, we determined the DEGs between HCs and SLE patients in each immune cell subset and selected those DEGs specific to immune cell subsets, such as B cells, that were common in the test and cohort data. By intersecting these DEGs with the eGenes, cell-type-specific key candidate genes and SNPs were identified.

### Hierarchical clustering and Factor analysis of type I IFN signaling-related signatures

To classify type I IFN signaling-related gene sets, the expression level of each of the 184 genes in this set was subjected to z-score normalization in each immune cell subset in SLE patients.

Using these expression data, we performed correlation-matrix-based hierarchical clustering. Factor analysis was used to determine whether the expression levels of multiple genes were regulated by a smaller number of unobserved continuous variables [17]. The Kaiser–Meyer–Olkin measure was used to verify the sampling adequacy of the analysis [18]. We applied the psych R package. The optimum number of factors was determined as under seven by a parallel analysis. After identifying the number of factors, oblique (promax; kappa = 4) rotation was used to obtain the final factor solution. Factor scores were calculated using Bartlett's method. The top 10 genes detected by factor analysis were used to calculate the signature score of each type I IFN signaling-related gene cluster.

**Correlations of clinical parameters with the OXPHOS and C6 gene signatures**

We excluded patients with an estimated glomerular filtration rate < 50 ml/min, because severe renal dysfunction might cause metabolic changes and affect the OXPHOS signature in PBMCs. The OXPHOS, C6, and ISRG scores in each immune cell subset were calculated by SVD. SLE patients were divided into the high and low signature score groups by Ward's hierarchical clustering method. After dividing the SLE patients with SDIs >0 into two groups according to the high/low OXPHOS or C6 signature score, enrichment analysis of each patient group to each clinical characteristic of SLE patients was performed. The signed −log10 *p*-values were visualized by heatmap. A *p*-value < 5% was considered significant.

**Reagents used for *in vitro* culture**

We used the B Cell Isolation Kit, mouse (Miltenyi Biotec) for isolation of splenic B cells and the Memory B Cell Isolation Kit II (Miltenyi Biotec) for isolation of human naive and memory B cells. The purity of the cells was confirmed to be > 95% in each experiment. Human B cells (1 x $10^5$/well) and mouse B cells (3 x $10^5$/well) were plated in 96-well U-bottom plates and cultured for 72 h with combinations of cytokines; a TLR9 agonist and IFN-α. The TLR9 agonists CpG ODN 2006 (Enzo Life Sciences) and ODN 1668 (Enzo Life Sciences) at 2.5 μg/ml were used for human and mouse TLR9 stimulation, respectively. Recombinant human IFN-α A (α 2b) (Pierce) and mouse IFN-α (PBL Assay Science) were used at a concentration of 1000 U/ml. Antibodies against the following proteins were used for flow cytometry: mouse CD3 (clone: 17A2), CD19 (clone: 1D3), CD138 (clone: 281-2), GL7 (clone: GL7), CD95 (Fas) (clone: Jo2) and B220 (clone: RA3-6B2), and human CD38 (clone: HIT2) and CD27 (clone: O323) (all from BioLegend). Antibodies against mouse CXCR5 and PD-1 (clone: J43) and human CD19 (clone: HIB19) were purchased from BD Biosciences. For ETC inhibition, human memory B cells (4 x $10^5$/well) were plated in 96-well U-bottom plates and then treated with the following ETC inhibitors (all from Sigma-Aldrich): 5 μM rotenone for mitochondrial complex I, 1 mM malonate for complex II, 1 μM antimycin A for complex III, and 1 mM KCN for complex IV.

**Transmission electron microscopy**

Human naive B cells, memory B cells, and PB cells were purified from four SLE patients and four healthy volunteers by magnetic cell sorting (Miltenyi Biotec). Splenic B cells were sorted from Prdx6 KO and WT littermate control mice by fluorescence activated cell sorting. The cells were fixed in Karnovsky's fixative (2% paraformaldehyde, 2.5% glutaraldehyde, and 0.08 M cacodylate buffer), dehydrated, and then embedded in epoxy resin. Tomography sections were imaged using the JEM-1011 (JEOL). Twenty cells per subset were analyzed, and mitochondria > 500 nm in diameter were defined as swollen [19]. We calculated the percentage of cells with swollen mitochondria.

**Extracellular flux analysis**

To investigate the metabolic status, we used a standardized protocol to measure the oxygen consumption rate and extracellular acidification rate using the Seahorse XF 96 analyzer as described previously [20]. In brief, $3 \times 10^5$ human B cells from PBMCs or $4 \times 10^5$ mouse splenic B cells/well were plated in 96-well Seahorse plates coated with Cell-Tak (Corning). As an indicator of mitochondrial function, the oxygen consumption rate was measured in real-time. B cells were treated sequentially with 1 μM oligomycin, 2 μM carbonyl cyanide-4-(trifluoromethoxy) phenylhydrazone (a mitochondrial uncoupler), and 0.5 μM rotenone plus 0.5 μM antimycin A (respiratory chain inhibitors).

**Immunization and quantification of NP-KLH-specific antibody responses**

Sex- and age-matched WT littermate control and Prdx6 KO mice were intraperitoneally administered 200 μg NP-KLH (Biosearch Technologies) emulsified with an equal amount of Imject Alum (Thermo Fisher Scientific). On day 8 following immunization, mouse sera were collected for ELISA, and isolated splenocytes were analyzed by flow cytometry. Anti-NP IgG and IgM levels were quantified by ELISA on plates coated with NP(4)-bovine serum albumin (Biosearch Technologies) as the capture antigen. Serially diluted pooled sera from NP-KLH-immunized C57BL/6 mice were utilized as a standard. Following incubation with sample and control sera, HRP-conjugated goat anti-mouse IgG or IgM (Bethyl Laboratories) and TMP substrate (KPL) were added to the plates.

**Statistical analysis**

Statistical tests were performed using Wilcoxon rank-sum test when sample sizes were more than five. Fisher's exact test was used to determine nonrandom associations between two categorical variables. The Boruta algorithm, a wrapper approach built around the random forest classification algorithm, was used for feature selection in SLE patients with neurologic disorders, using perc = 99 as the threshold. Boruta is an open-source package for Python available at https://github.com/scikit-learn-contrib/boruta_py.
