## Supplementary Table for "Immune cell multi-omics analysis reveals contribution of oxidative phosphorylation to B cell functions and organ damage of lupus"

**Online supplementary materials and methods**

**Online Supplemental Table S1. Each ETC complex gene list used in this study.**

| Complex I | Complex II | Complex III | Complex IV | Complex V |
|---|---|---|---|---|
| NDUFA1 | SDHA | CYC1 | COX4I1 | ATP5F1A |
| NDUFA10 | SDHB | UQCR10 | COX4I2 | ATP5F1B |
| NDUFA11 | SDHC | UQCR11 | COX5A | ATP5F1C |
| NDUFA12 | SDHD | UQCRB | COX5B | ATP5F1D |
| NDUFA13 | | UQCRC1 | COX6A1 | ATP5F1E |
| NDUFA2 | | UQCRC2 | COX6A2 | ATP5IF1 |
| NDUFA3 | | UQCRFS1 | COX6B1 | ATP5MC1 |
| NDUFA5 | | UQCRH | COX6B2 | ATP5MC2 |
| NDUFA6 | | UQCRQ | COX6C | ATP5MC3 |
| NDUFA7 | | | COX7A1 | ATP5MD |
| NDUFA8 | | | COX7A2 | ATP5ME |
| NDUFA9 | | | COX7B | ATP5MF |
| NDUFAB1 | | | COX7B2 | ATP5MG |
| NDUFB1 | | | COX7C | ATP5MPL |
| NDUFB10 | | | COX8A | ATP5PB |
| NDUFB11 | | | COX8C | ATP5PD |
| NDUFB2 | | | | ATP5PF |
| NDUFB3 | | | | ATP5PO |
| NDUFB4 | | | | |
| NDUFB5 | | | | |
| NDUFB6 | | | | |
| NDUFB7 | | | | |
| NDUFB8 | | | | |
| NDUFB9 | | | | |
| NDUFC1 | | | | |
| NDUFC2 | | | | |
| NDUFS1 | | | | |
| NDUFS2 | | | | |

| |
|---|
| NDUFS3<br>NDUFS4<br>NDUFS5<br>NDUFS6<br>NDUFS7<br>NDUFS8<br>NDUFV1<br>NDUFV2<br>NDUFV3 |

1

**Online Supplemental Table S2. Results of cis-eQTL analysis using RNA-seq and genotyping data from our IMD patients and HCs.**

| SLE GWAS Top | eQTL Top | Subset | $r^2$_JPN | $r^2$_EUR | eGene | Aliases and/or Function of eGene | References |
|---|---|---|---|---|---|---|---|
| rs1385374 | rs3765107 | DN B | 0.955 | 1.000 | SLC15A4 | Histidine transporter on lysosomes | SLC15A4 in B cells regulates type I IFN production through regulation of TLR7/9 as well as mTORC1 signaling [1]. |
| rs1385374 | rs3765107 | Naïve B | 0.955 | 1.000 | SLC15A4 | | |
| rs11059927 | rs3765107 | DN B | 0.955 | 1.000 | SLC15A4 | | |
| rs11059927 | rs3765107 | Naive B | 0.955 | 1.000 | SLC15A4 | | |
| rs2422345 | rs11484635 | USM B | 0.995 | 0.995 | TNFSF4 | OX40L | OX40L-expressing B cells supports plasma cell differentiation and Tfh maturation [2]. |
| rs4916342 | rs11484635 | USM B | 0.995 | 0.930 | TNFSF4 | | |
| rs10753074 | rs11484635 | USM B | 0.995 | 0.991 | TNFSF4 | | |
| rs4426778 | rs7686702 | Naive B | 1.000 | 0.985 | BANK1 | B cell scaffold protein with ankyrin repeats 1 | BANK1 on B cell promotes antigen presentation ability and autoantibody production [3]. |
| rs11085725 | rs11085727 | SM B | 0.962 | 1.000 | TYK2 | Tyrosine kinase 2 | |

| | | | | | | | |
|---|---|---|---|---|---|---|---|
| rs11085725 | rs35251378 | DN B | 0.991 | 0.995 | TYK2 | | Tyk2 tyrosine kinase expression Is required for the maintenance of mitochondrial respiration in primary pro-B lymphocytes [4]. |
| rs11085725 | rs35251378 | Plasmablast | 0.991 | 0.995 | TYK2 | | |
| rs11085727 | rs11085727 | SM B | 0.961 | 0.995 | TYK2 | | |
| rs11085727 | rs35251378 | DN B | 0.961 | 0.995 | TYK2 | | |
| rs11085727 | rs35251378 | Plasmablast | 0.961 | 0.995 | TYK2 | | |
| rs1170426 | rs1728797 | USM B | 0.992 | 1.000 | ZFP90 | Zink finger protein | ZFP90 transgenic mice showed altered expression of OXPHOS related genes [5]. |
| rs1170426 | rs4783650 | Naive B | 1.000 | 0.849 | ZFP90 | | |
| rs131658 | rs5754467 | SM B | 0.956 | 0.800 | UBE2L3 | Ubiquitin conjugating enzyme E2L3 | UBE2N, UBE2L3, and UBE2D2/3 contribute to Parkin-mediated mitophagy [6]. |
| rs3747093 | rs5754467 | SM B | 0.996 | 0.974 | UBE2L3 | | |
| rs2061831 | rs2061831 | Naive B | 0.995 | 1.000 | BLK | B lymphoid tyrosine kinase | BLK and BANK1 act through PLC gamma 2 in B cell signaling [7]. |
| rs2061831 | rs2409780 | USM B | 0.995 | 1.000 | BLK | | |
| rs2254546 | rs2254546 | Plasmablast | 0.937 | 0.993 | BLK | | |
| rs2618444 | rs2061831 | Naive B | 0.990 | 1.000 | BLK | | |
| rs2618444 | rs2409780 | USM B | 0.995 | 1.000 | BLK | | |
| rs2618476 | rs2061831 | Naive B | 0.924 | 0.950 | BLK | | |
| rs2618476 | rs2409780 | USM B | 0.919 | 0.950 | BLK | | |

| | | | | | | | |
|---|---|---|---|---|---|---|---|
| rs2736332 | rs2061831 | Naive B | 0.995 | 0.876 | BLK | | |
| rs2736332 | rs2409780 | USM B | 1.000 | 0.876 | BLK | | |
| rs2736337 | rs2061831 | Naive B | 0.962 | 0.995 | BLK | | |
| rs2736337 | rs2409780 | USM B | 0.957 | 0.995 | BLK | | |
| rs2736340 | rs2061831 | Naive B | 0.957 | 0.995 | BLK | | |
| rs2736340 | rs2409780 | USM B | 0.952 | 0.995 | BLK | | |
| rs7812879 | rs2254546 | Plasmablast | 0.911 | 0.986 | BLK | | |
| rs13277113 | rs2061831 | Naive B | 0.897 | 0.990 | BLK | | |
| rs13277113 | rs2409780 | USM B | 0.893 | 0.990 | BLK | | |
| rs2299864 | rs1322178 | Plasmablast | 0.981 | 0.964 | ATG5 | Autophagy related 5 | Mitochondrial quality control after oxidative damage. Localizes to punctae on mitochondria [8]. |
| rs9373839 | rs1322178 | Plasmablast | 0.981 | 0.959 | ATG5 | | |
| rs2732549 | rs146368551 | Naive B | 0.821 | 0.913 | CD44 | Cell surface glycoprotein | CD44lo to CD44hi conversion of esophageal keratinocytes induces mitochondrial dysfunction and OXPHOS [9]. |
| rs3794060 | rs732934 | Naive B | 1.000 | 1.000 | NADSYN1 | NAD synthetase 1 | |
| rs3794060 | rs732934 | USM B | 1.000 | 1.000 | NADSYN1 | | |

| | | | | | | | |
|---|---|---|---|---|---|---|---|
| rs6445975 | rs4681845 | Naive B | 0.914 | 0.971 | PXK | PX domain containing serine/theronine kinase like | PXK influenced the rate of BCR internalization [10]. |
| rs6445975 | rs4681845 | SM B | 0.914 | 0.971 | PXK | | |
| rs6445975 | rs4681845 | USM B | 0.914 | 0.971 | PXK | | |
| rs7258015 | rs2304237 | Naive B | 1.000 | 0.979 | ICAM3 | Ig-like adhesion molecule | ICAM-3 crosslinking induced the production of reactive oxygen species (ROS), which are known to be involved in the control of endothelial cell-cell contacts [11]. |
| rs7258015 | rs2304237 | USM B | 1.000 | 0.979 | ICAM3 | | |
| rs73366469 | rs73366469 | Plasmablast | 0.853 | 1.000 | NCF1 | Subunit of NADPH oxidase | |
| rs9782955 | rs2104125 | DN B | 0.990 | 0.959 | LYST | Lysosomal trafficking regulator | Lyst mutation results in elevated levels of oxidative damage to lipid membranes [12]. |
| rs9782955 | rs2104125 | Naive B | 0.990 | 0.959 | LYST | | |
| rs9782955 | rs2104125 | SM B | 0.990 | 0.959 | LYST | | |

**Online Supplemental Table S3. Parent subset common DEGs in test and validation cohorts with the information of *cis*-eQTL analysis**

| Gene | subset common DEGs | | | | | | | | eQTL & GWAS | | | | |
|---|---|---|---|---|---|---|---|---|---|---|---|---|---|
| | B | | CD4 | | DC | | Mono | | B | CD4 | DC | Mono | others |
| | test | valida-tion | test | valida-tion | test | valida-tion | test | valida-tion | | | | | |
| CLSPN | | | | | | | | | | | | | |
| GCA | | | | | | | | | | | | | |
| IL18R1 | | | * | | | | | | | * | | | |
| IL18RAP | | | | | | | | | | | | | |
| IL7 | * | | * | | | | | | * | * | | | |
| LGALS9 | * | | | | * | | * | | * | | * | * | |
| NCF2 | | | | | | | | | | | | | |
| NEU1 | | | | | | | | | | | | | |
| PDHX | | | | | | | | | | | | | |
| PHKG2 | | | | | | | | | | | | | |
| PKIA | * | | * | | | | | | * | * | | | |
| PRDM1 | | | | | | | | | | | | | |
| PRDX6 | * | | | | | | | | * | | | | |
| PRNP | | | | | | | | | | | | | |
| PYCARD | | | * | | | | | | | * | | | |
| RGS1 | | | * | | | | | | | * | | | |
| RPL10A | | | | | | | * | | | | | * | |
| SERBP1 | | | | | | | | | | | | | |

An asterisk (*) indicates a candidate gene which was picked up as a DEG in both cohorts and also eGene from eQTL and GWAS.

**Online Supplemental Table S4. Gene lists of each type I IFN signaling-related gene cluster.**

| C1 | C2 | C3 | C4 | C5 | C6 |
|---|---|---|---|---|---|
| RNF135 | ITCH | POLR1D | TRIM25 | RIPK2 | POLR3H |
| MB21D1 | TRAIP | PCBP2 | ADAR | PTPN22 | HSP90AB1 |
| IKBKE | IRF4 | RPS27A | STAT2 | TBK1 | HSPD1 |
| PTPN6 | DHX9 | UBA52 | DHX58 | PTPN2 | XRCC5 |
| UBC | SAMHD1 | TRAF3IP1 | OAS3 | POLR2K | XRCC6 |
| CDC37 | CTNNB1 | PRKDC | IRF7 | POLR3GL | FADD |
| UBA7 | ZC3HAV1 | CRCP | OAS2 | TAX1BP1 | IRF2 |
| IRF1 | POLR3B | CREBBP | C19orf66 | UBE2K | GBP2 |
| RELA | EP300 | RNF216 | IRF9 | MRE11A | HAVCR2 |
| HLA-A | POLR3A | PTPN11 | TRIM38 | IFNAR1 | PYCARD |
| MYD88 | ZBTB20 | LRRFIP1 | DDX58 | IFNAR2 | PIN1 |
| TNFAIP3 | JAK1 | DHX36 | IFIH1 | | POLR2L |
| IRF5 | REL | PPM1B | STAT1 | | POLR2F |
| NLRC5 | IRF8 | TRAF3 | SP100 | | UBB |
| STAT6 | RNASEL | OTUD5 | IFI16 | | HMGB2 |
| TYK2 | POLR3F | RELB | NMI | | ISG20 |
| PTPN1 | TRIM32 | NFKB1 | UBE2L6 | | HMGB1 |
| TICAM1 | LSM14A | NFKB2 | BST2 | | POLR3K |
| NLRX1 | CYLD | IRAK1 | IFI35 | | POLR1C |
| TRIM56 | DDX3X | PLCG2 | MX2 | | POLR3C |
| POLR3E | | POLR2H | OASL | | IP6K2 |
| MAVS | | POLR3D | HERC5 | | HLA-E |
| TIRAP | | | XAF1 | | HLA-F |
| HLA-B | | | IFIT1 | | |
| HLA-C | | | IFIT3 | | |
| HLA-H | | | IFITM1 | | |
| MUL1 | | | IFI6 | | |
| TMEM173 | | | IFI27 | | |
| DDX41 | | | MX1 | | |
| SHMT2 | | | RSAD2 | | |
| HLA-G | | | OAS1 | | |
| FLOT1 | | | ISG15 | | |
| IRF3 | | | USP18 | | |

| | | | |
|---|---|---|---|
| POLR2E | | | IFITM2 |
| | | | IFITM3 |
| | | | TREX1 |
| | | | PSMB8 |
| | | | TRIM21 |

**Online Supplemental Table S5. Correlation of C1 to C5 signatures with the OXPHOS signature in each immune cell subset**

Test cohort

| | **Cluster1** | | | **Cluster2** | | | **Cluster3** | | | **Cluster4** | | | **Cluster5** | | |
|---|---|---|---|---|---|---|---|---|---|---|---|---|---|---|---|
| **subset** | *r* | *q* | | *r* | *q* | | *r* | *q* | | *r* | *q* | | *r* | *q* | |
| Naive B | -0.34 | 0.03 | * | -0.24 | 0.11 | | -0.21 | 0.24 | | 0.3 | 0.09 | | 0.33 | 0.05 | * |
| SM B | 0.03 | 0.85 | | -0.63 | 1.3 xE-5 | **** | 0.47 | 7.2 xE-3 | ** | 0.53 | 5.1 xE-4 | *** | 0.31 | 0.05 | * |
| USM B | -0.11 | 0.49 | | -0.46 | 3.8 xE-3 | ** | 0.29 | 0.12 | | 0.3 | 0.09 | | 0.41 | 0.03 | * |
| DN B | -0.44 | 2.9 xE-3 | ** | -0.6 | 3.9 xE-5 | **** | -0.08 | 0.7 | | 0.02 | 0.91 | | 0.24 | 0.11 | |
| Plasma-blast | -0.5 | 4.8 xE-4 | *** | -0.5 | 9.0 xE-4 | *** | -0.23 | 0.21 | | 0.45 | 6.4 xE-3 | ** | 0.37 | 0.03 | * |
| Th1 | -0.51 | 3.3 xE-4 | *** | -0.51 | 5.4 xE-4 | *** | -1.7 xE-3 | 0.99 | | 0.66 | 4.8 xE-6 | ***** | 0.28 | 0.07 | |
| Th2 | -0.59 | 4.9 xE-5 | **** | -0.24 | 0.11 | | -0.22 | 0.21 | | 0.29 | 0.09 | | 0.27 | 0.08 | |
| Th17 | -0.56 | 7.6 xE-5 | **** | -0.36 | 0.01 | * | -0.22 | 0.21 | | 0.24 | 0.15 | | 0.45 | 8.3 xE-3 | ** |
| Tfh | -0.57 | 7.6 xE-5 | **** | -0.37 | 0.01 | * | -0.37 | 0.05 | * | 0.32 | 0.07 | | 0.35 | 0.04 | * |
| Naive CD4 | -0.42 | 4.3 xE-3 | ** | -0.37 | 0.01 | * | -0.28 | 0.12 | | 0.25 | 0.14 | | 0.38 | 0.03 | * |
| Mem CD4 | -0.7 | 3.2 xE-7 | ****** | -0.16 | 0.26 | | -0.27 | 0.12 | | 0.29 | 0.09 | | 0.51 | 3.4 xE-3 | ** |
| Fr. II eTreg | -0.58 | 5.1 xE-5 | **** | -0.39 | 8.1 xE-3 | ** | -0.33 | 0.07 | | 0.07 | 0.68 | | 0.46 | 8.3 xE-3 | ** |
| Naïve CD8 | -0.43 | 2.9 xE-3 | ** | -0.38 | 9.8 xE-3 | ** | -0.17 | 0.32 | | 0.33 | 0.07 | | 0.1 | 0.53 | |

| | | | | | | | | | | | | | | | |
|---|---|---|---|---|---|---|---|---|---|---|---|---|---|---|---|
| Mem CD8 | -0.35 | 0.02 | * | -0.52 | 5.3 xE-4 | *** | -0.07 | 0.7 | | 0.64 | 5.5 xE-6 | ***** | 0.33 | 0.05 | * |
| mDC | -0.56 | 7.6 xE-05 | **** | -0.59 | 3.9 xE-05 | **** | -0.41 | 0.02 | * | 0.13 | 0.43 | | 0.31 | 0.05 | * |
| pDC | -0.38 | 9.6 xE-03 | ** | -0.43 | 3.8 xE-03 | ** | -0.11 | 0.57 | | 0.21 | 0.18 | | 0.3 | 0.05 | * |
| CD16p Mono | -0.63 | 1.2 xE-05 | **** | -0.64 | 1.3 xE-05 | **** | -0.49 | 6.8 xE-03 | ** | -0.12 | 0.45 | | 0.19 | 0.21 | |
| CD16n Mono | -0.58 | 4.9 xE-05 | **** | -0.4 | 8.1 xE-03 | ** | -0.28 | 0.12 | | 0.22 | 0.18 | | 0.37 | 0.03 | * |
| NK | -0.34 | 0.02 | * | -0.47 | 1.6 xE-03 | ** | 0.05 | 0.77 | | 0.35 | 0.06 | | -0.07 | 0.64 | |

**Online Supplemental Table S6. Clinical characteristics of the enrolled SLE patients**

| **Characteristics** | **HC** | | **SLE** | |
|---|---|---|---|---|
| | **Test cohort** | **Validation cohort** | **Test cohort** | **Validation cohort** |
| | n = 37 | n = 55 | n = 49 | n = 58 |
| **Age** | 52.2 ± 13.8 | 44.2 ± 15.0 | 49.3 ± 10.6 | 45.9 ± 13.3 |
| **Gender, female** | 32 (86.5%) | 37 (67.3%) | 44 (89.8%) | 50 (86.2%) |
| **Disease duration** | NA | NA | 19.5 ± 12.0 | 13.2 ± 10.8 |
| **SLEDAI-2K** | NA | NA | 5.2 ± 3.5 | 6.9 ± 5.2 |
| **SDI** | NA | NA | 1.0 ± 1.2 | 0.9 ± 1.4 |
| **Complement decrease** | NA | NA | 22 (44.9%) | 28 (48.3%) |
| **Leukopenia** | NA | NA | 41 (85.4%) | 46 (80.7%) |
| **Lymphopenia** | NA | NA | 35 (71.4%) | 44 (77.2%) |
| **Plasmablast (% in total B)** | 1.4 ± 1.6 | 2.6 ± 2.7 | 9.0 ± 8.9 | 9.9 ± 9.8 |
| **Anti-dsDNA positive** | NA | NA | 16 (32.7%) | 22 (38.6%) |
| **Anti-dsDNA titer** | NA | NA | 21.2 ± 31.9 | 24.7 ± 60.2 |
| **Anti-Sm positive** | NA | NA | 3 (6.1%) | 9 (16.4%) |
| **Anti-RNP positive** | NA | NA | 14 (28.6%) | 20 (37.0%) |
| **APA positive** | NA | NA | 16 (32.7%) | 19 (36.5%) |
| **Daily PSL dose (mg/day)** | NA | NA | 5.4±2.4 | 7.5±10.4 |
| **PSL** | NA | NA | 46 (93.9%) | 49 (84.5%) |
| **Azathioprine** | NA | NA | 3 (6.1%) | 7 (12.1%) |
| **Cyclophosphamide (p.o.)** | NA | NA | 1 (2.0%) | 0 (0%) |
| **Cyclosporin A** | NA | NA | 6 (12.2%) | 6 (10.3%) |
| **Tacrolimus** | NA | NA | 8 (16.3%) | 6 (10.3%) |
| **Mycophenolate mofetil** | NA | NA | 2 (4.1%) | 9 (15.6%) |

**Table S7. Sorting Panel of Each Immune Cell Subset**

| Parent | Subset Name | Gating Strategy | Comment |
|---|---|---|---|
| B cell | Naïve B | CD3-CD19+IgD+CD27- | |
| | Switched memory B (SM B) | CD3-CD19+IgD-CD27+ | test cohort only |
| | Switched memory B (SM B) | CD3-CD19+IgD-CD27+CD38- | validation cohort only |
| | Unswitched memory B (USM B) | CD3-CD19+IgD+CD27+ | |
| | Double Negative B (DN B) | CD3-CD19+IgD-CD27- | |
| | Plasmablast | CD3-CD19+IgD-CD27++CD38+ | |
| T cell | Th1 | CD3+CD4+CD25-CD45RA-CXCR5-CCR5-CXCR3+ | test cohort only |
| | Th1 | CD3+CD4+CD8-CD25-CCR4-CCR6-CXCR3+CXCR5- in non-NaiveCD4 | validation cohort only |
| | Th2 | CD3+CD4+CD25-CD45RA-CXCR5-CCR5-CXCR3- | test cohort only |
| | Th2 | CD3+CD4+CD8-CD25-CCR4+CCR6-CXCR3-CXCR5- in non-NaiveCD4 | validation cohort only |
| | Th17 | CD3+CD4+CD25-CD45RA-CXCR5-CCR5+CXCR3- | test cohort only |
| | Th17 | CD3+CD4+CD8-CD25-CCR6+CXCR3-CXCR5- in non-NaiveCD4 | validation cohort only |
| | Tfh | CD3+CD4+CD25-CXCR5+ | test cohort only |
| | Tfh | CD3+CD4+CD8-CD25-CXCR5+ in non-NaiveCD4 | validation cohort only |
| | Naïve CD4 T (Naïve CD4) | CD3+CD4+CD45RA+ | test cohort only |
| | Naïve CD4 T (Naïve CD4) | CD3+CD4+CD8-CCR7+CD45RA+ | validation cohort only |
| | Memory CD4 T (Mem CD4) | CD3+CD4+CD25-CD45RA- | test cohort only |
| | Memory CD4 T (Mem CD4) | CD3+CD4+CD8-CD25- in non-NaiveCD4 | validation cohort only |
| | Fraction II effector regulatory T (Fr. II eTreg) | CD3+CD4+CD25highCD45RA- | test cohort only |

| | | | |
|---|---|---|---|
| | Fraction II effector regulatory T (Fr. II eTreg) | CD3+CD4+CD8-CD25highCD45RA- | validation cohort only |
| | Naïve CD8 T (Naïve CD8) | CD3+CD19-CD8+CD45RA+ | test cohort only |
| | Naïve CD8 T (Naïve CD8) | CD3+CD4-CD8+CD45RA+CCR7+ | validation cohort only |
| | Memory CD8 T (Mem CD8) | CD3+CD19-CD8+CD45RA- | test cohort only |
| | NK | CD3-CD19-CD56+CD14- | test cohort only |
| Monocyte | CD16 positive monocytes (CD16p Mono) | CD3-CD19-CD56-HLADR+CD14+CD16+ | |
| | CD16 negative monocytes (CD16n Mono) | CD3-CD19-CD56-HLADR+CD14+CD16- | |
| Dendritic Cell | Myeloid Dendritic Cells (mDC) | CD3-CD19-CD56-HLADR+CD14-(CD16-)CD11c+CD123- | |
| | Plasmacytoid Dendritic Cells (pDC) | CD3-CD19-CD56-HLADR+CD14-(CD16-)CD11c-CD123+ | |
